# A unified theory for the computational and mechanistic origins of grid cells

**DOI:** 10.1101/2020.12.29.424583

**Authors:** Ben Sorscher, Gabriel C. Mel, Samuel A. Ocko, Lisa Giocomo, Surya Ganguli

## Abstract

The discovery of entorhinal grid cells has generated considerable interest in how and why hexagonal firing fields might mechanistically emerge in a generic manner from neural circuits, and what their computational significance might be. Here we forge an intimate link between the computational problem of path-integration and the existence of hexagonal grids, by demonstrating that such grids arise generically in biologically plausible neural networks trained to path integrate. Moreover, we develop a unifying theory for why hexagonal grids are so ubiquitous in path-integrator circuits. Such trained networks also yield powerful mechanistic hypotheses, exhibiting realistic levels of biological variability not captured by hand-designed models. We furthermore develop methods to analyze the connectome and activity maps of our trained networks to elucidate fundamental mechanisms underlying path integration. These methods provide an instructive roadmap to go from connectomic and physiological measurements to conceptual understanding in a manner that might be generalizable to other settings.

## 1 Introduction

The history of neuroscience is punctuated by striking experimental observations of single cell tuning properties in different brain regions that appear almost universally across many different species (e.g. center-surround receptive fields in the retina [1], Gabor-like receptive fields in primary visual cortex [2], and temporal wavelet like receptive fields in primary auditory cortex [3]). More recently, the experimental observation of another striking and ubiquitous single cell tuning property has engendered intense interest in neuroscience: the regular hexagonal firing patterns across space of grid cells in the medial entorhinal cortex (MEC). These patterns have been observed as a function of spatial position in mice [4], rats [5], and bats [6] and as a function of gaze position in monkeys [7]. Also fMRI studies have revealed evidence for grid like representations in humans [8]. The regularity and ubiquity of grid cells raise two distinct but related classes of scientific questions. First, what circuit mechanisms might give rise to grid cells? In particular, how might hexagonal grid cell firing patterns arise in a *generic* manner from neural circuit connectivity and dynamics? And second, what computational reasons might explain the seemingly universal existence of grid cells across many branches of the phylogenetic tree? In essence, if hexagonal grid cells are evolution’s answer to an ethologically relevant computational question, then what is that question?

On the mechanistic side, many works have focused on hand-tuning the connectivity of model recurrent neural circuits with a center-surround structure specifically to generate grid cell firing patterns [9–12], building on prior models of head-direction cells [13–17] and place cells [18–20]. Such continuous attractor models can robustly integrate and store 2D positional information via path integration [11]. More recent attractor networks that incorporate plastic inputs from landmark cells can explain why grid cells deform in irregular environments [12, 13], and when they either phase shift or remap in altered virtual reality environments [21]. However, such models designed through hand-tuning by theorists raise two questions. First, such models involve many choices about circuit connectivity and dynamics, and it is unclear how generic such choices are. In essence could there be vastly more different classes of attractor neural networks that both path-integrate and generate hexagonal firing patterns as a result? Second, none of these recurrent network models show that grid-like firing patterns are *required* to solve navigational tasks optimally. Thus they cannot demonstrate that hexagonal firing patterns naturally arise as the solution to a computational problem, precisely because the hexagonal patterns are simply assumed in the first place by hand-tuning the recurrent connectivity of these networks.

On the other hand, normative models attempt to shed light on the question of *why* grid firing patterns might be found in many species by demonstrating that these patterns are optimal for a solving a particular task. For example, [22] demonstrated that single neurons that receive place cell inputs through plastic synaptic weights undergoing Oja’s learning rule develop grid like receptive fields (RFs) with a square lattice structure. If these synapses are also constrained to be positive, then these same neurons learn hexagonal grid-like RFs. Since Oja’s learning rule, without weight constraints, performs principal components analysis (PCA) [23], this work essentially demonstrates that grid like RFs can form an efficient representation of place cell *input* patterns. An alternative theoretical approach focuses on optimal representations of the statistics of possible transitions in a spatial environment [24, 25], finding that square grid representations are optimal in square environments. Other works have assumed the existence of grid like representations with different lattice structures, finding hexagonal lattices outperform other lattice structures in terms of either decoding position under noise [26], or a notion of economy [27]. However, unlike the hand-tuned continuous attractor network models described above, these normative approaches primarily tackle the issue of spatial representations, and do not tackle the central issue of how neural circuits might actually solve the problem of path integration, which is thought to be a primary computational function of entorhinal cortex, whether in real space, or in abstract spaces [28, 29].

More recent normative approaches have tackled this central issue head on by training, rather than hand-tuning, neural networks to accurately solve the navigational problem of path integration. Indeed, [30] found that square grid cells spontaneously emerge in square environments in such trained networks. Also [31] suggested hexagonal grid cells emerge even in square environments in such trained networks, though the grid cell patterns were highly heterogeneous. The process of training neural networks to solve different computational problems can be a powerful method for hypothesis generation in neuroscience, yielding a more unbiased search process in the space of circuit solutions than might be possible through the imagination of theorists alone. Indeed the representations of trained rather than hand-designed networks have been successfully compared to actual neural representations in the retina [12, 32–34], inferotemporal cortex, V4, and V1 [35, 36], motor cortex [37] and prefrontal cortex [38]. However in comparisons to the behavior of entorhinal representations, the trained networks in [30] and [31] exhibit mismatches along two important dimensions. First, as we will show below, the learned lattice structure is inconsistent with data (square versus hexagonal in the case of [30], and smooth and random versus truly hexagonal in the case of [31]). Second, as we will see below, the methods of training neural networks in [31] in a fixed environment do not yield grid cells whose representations generalize to expanded environments.

Despite these mismatches, [30, 31] exemplify interesting advances in generating, in a more unbiased manner, neural networks that path-integrate with grid like representations, and they naturally suggest important, yet unanswered questions. First, when and why do grid like representations spontaneously emerge from neural circuits that are trained to solve navigational problems, or other normative problems like efficient spatial encoding [22]? And, second, if grid-like representations appear, when and why are they sometimes square, hexagonal, or heterogeneous? Third, focusing on the networks trained to path-integrate, what circuit mechanisms yield grid like responses? Are these circuit mechanisms for both path-integration and grid-cells in trained models at all related to the circuit mechanisms dreamed up by theorists in prior hand-tuned models? In essence, how can we obtain conceptual insight into how circuit connectivity and dynamics conspire to yield the emergent computational functions of these trained circuits? This latter question is a foundational question for neuroscience in general, especially as we encounter more and more circuit models generated via machine-learning based training methods [34]. In such models we have access to the *entire* connectome and the activity patterns of *every* neuron. Thus the process by which we might go from such connectivity and activity data to conceptual understanding could be highly instructive in teaching us how we might leverage data generated by expensive investments in measuring entire connectomes [39] and very large scale brain activity maps [40]. In essence, methods of understanding complete synaptic and neural data from computational models could be informative in elucidating what conceptual understanding we could potentially extract from large scale connectomics and activity data in other settings [41], and moreover, *how* we can efficiently extract such understanding.

In this work, we address all the questions raised above concerning relations between the computational and mechanistic origins of grid cells across normative representational models, and hand-tuned as well as trained path integrators. First, we obtain more robust hexagonal grid like representations in neural networks trained to path integrate by introducing a simple biological constraint on firing rates, namely that they are non-negative. Second, we train them across multiple environments to develop neural networks that not only path-integrate with hexagonal grid cells in a square environment, but also generalize this hexagonal pattern outside the original square, as this environment is expanded. Our networks thus go beyond prior trained networks, which do not yield robust hexagonal grids or generalize in expanded environments. Third, we develop a general unified theory for *why* grid cells can spontaneously emerge in diverse normative as well as mechanistic models, including in recurrent networks trained to path-integrate [30, 31] as well as feedforward networks trained to efficiently encode place cell inputs [22]. Fourth, across all these works, our general theory explains why and when different grid lattice structures (i.e. square, hexagonal, and heterogeneous) spontaneously emerge. Fifth, we develop novel algorithmic methods to extract, from the seemingly highly unstructured connectomes of such trained networks, a conceptual understanding of how such networks both path integrate as well as mechanistically generate hexagonal grid cell responses. These methods thus open the black box of the network’s connectome and activity maps, and lay bare conceptually interpretable mechanistic hypotheses for entorhinal circuit operation, including hidden connectivity patterns mediating path integration. Sixth, we relate our conceptual understanding of the circuit mechanisms underlying trained grid-cell path integrators to those of hand-tuned path integrators, showing that the former are interesting and much more general versions of the latter. Thus overall, we hope this work provides broad theoretical clarity on potential entorhinal function across several dimensions, provides a unified theory of neural network based path-integration and the spontaneous emergence of grid cell like structures, and provides an instructive road map to go from large scale connectomic and activity data to conceptual understanding, a road map that might be generalized to other settings.

## 2 Results

### 2.1 Diverse lattice structures and generalization properties of neural networks trained to path integrate

Path integration refers to the process of integrating instantaneous velocity signals to obtain an estimate of current position, and is thought to be a computational function mediated by grid cells in MEC. We therefore sought to elucidate the conditions under which the computational demand of path integration alone might naturally lead to the spontaneous emergence of hexagonal grid cells in a trained neural network. In particular, we simulated an animal exploring a square environment following a smooth random walk (Figure 1A). As the animal moves, different subsets of simulated place cells become active (place cell receptive field centers are shown as dots in Figure 1A). At each time step, the network receives the animal’s 2-dimensional velocity 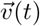 as input. The velocity signal is integrated by the network’s recurrently connected units, and the entire network is trained to report its current position by creating as output the firing rates of a set of simulated place cells. Figure 1B shows a schematic for neural network models of this type.

**Figure 1:**
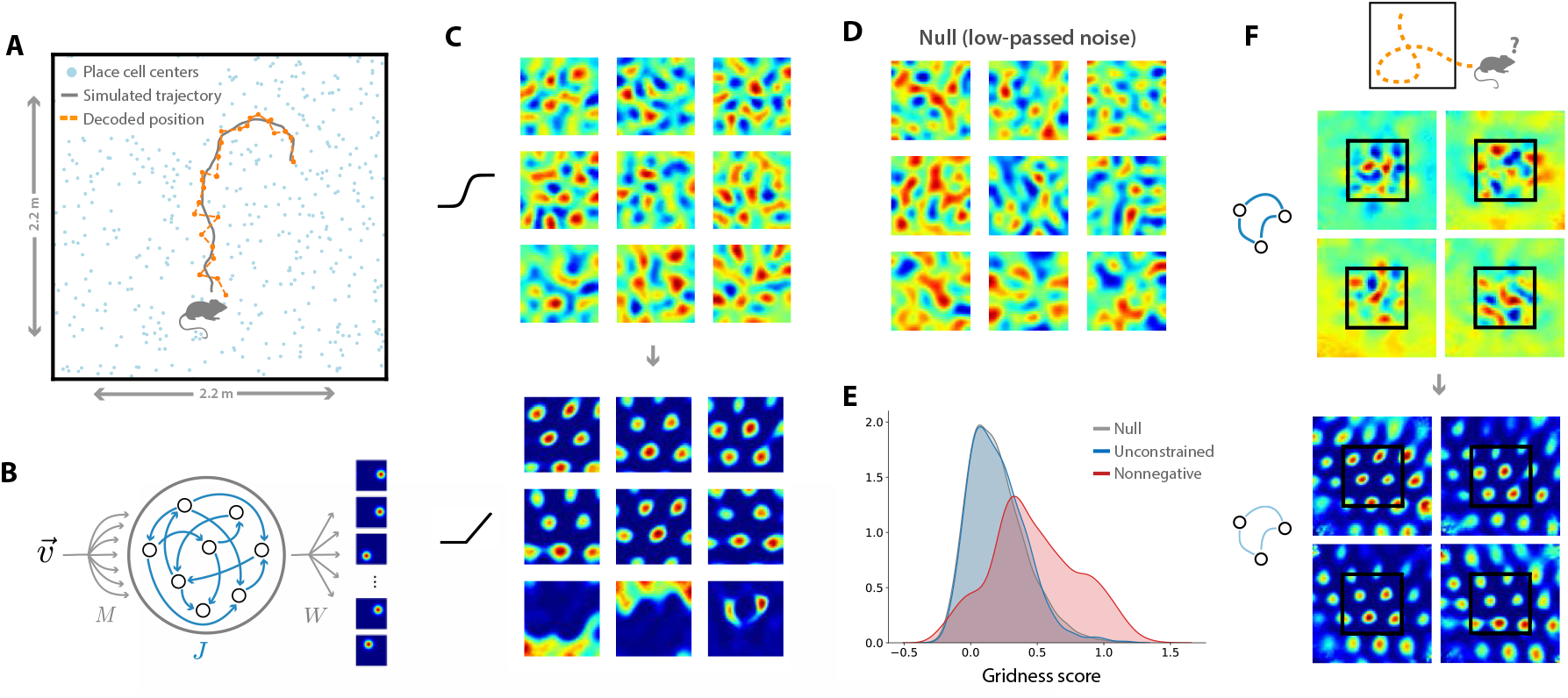
Learned spatial representations of neural networks trained to path integrate. (A) A simulated animal trajectory (grey curve), and the decoded position from the network’s output (orange curve). Place cell centers (blue dots) are distributed uniformly and isotropically over a square enclosure. (B) Our general model architecture includes a velocity input which is fed to a set of recurrently connected hidden units. These hidden units must then generate a desired place cell output representation. (C) A nonnegativity constraint on firing rates induces hexagonal grids. Learned hidden unit representations when the single neuron nonlinearity is sigmoidal (top) and rectified linear (ReLu) (bottom). (D) A null model of single neuron spatial representations obtained by low-pass filtering spatial noise (see Methods 4.3). (E) A distribution of grid scores for all hidden units in the sigmoidal network with unconstrained firing rates in panel C top (blue), the null model in panel D (grey), and the ReLU network with nonnegative firing rates in panel C bottom. The null distribution of grid scores closely matches that of the unconstrained network, and the nonnegative network exhibits significantly higher grid scores. (E) Generalizing beyond the training environment. Without weight decay, the RNN freezes at the boundary of the training environment (top). With weight decay, the RNN continues path integrating outside the training environment (bottom). See Methods 4.2 for all details of the training procedure.

We began by reproducing the results of [31], replicating their specific network architecture and path integration task (see Methods 4.2 for training details). After training, neurons in the network developed firing fields that exhibited clear grid-like tuning. However, we found that networks trained as above had two biologically unrealistic characteristics. First, the spatial tuning was highly heterogeneous (Figure 1C, upper), qualitatively unlike the regular hexagonal tuning of many entorhinal grid cells. Indeed the learned maps did not have higher grid scores [42] than those drawn from a null distribution of low-pass-filtered noise (Figure 1D,E, and Methods 4.3). Second, the network was unable to path integrate outside of the training arena (Figure 1F, upper). When the animal’s simulated walk was chosen to pass beyond a removed wall of the square environment used during training, the hidden unit activities in the trained network usually froze at their boundary values, rather than continuing to fire in a spatially informative way, as biological grid cells do [43]).

We found that we could obtain similar results in a significantly simpler recurrent neural network (RNN) architecture (shown in Figure 1B), without all the complexity of the architecture used in [31] (for training details, see Methods 4.2). The simple RNN learned to path integrate comparably well, and developed qualitatively similar grid-like patterns in its hidden units. The simple RNN architecture has a couple of key advantages: (1) the grid cells are recurrently connected, like grid cells in MEC, and unlike those in [31], and (2) this architecture corresponds exactly to traditional path integrator models of grid cells [9–20], except that while the recurrent weights in traditional models are chosen by hand, ours are learned, likely in a less biased fashion, over the course of training.

Next, we found that two simple changes to the training procedure encouraged the network to (1) reliably learn regular hexagonal grids like those in MEC, and (2) learn a path integration mechanism which generalizes outside of the training environment, resolving both of the problems identified in the top panels of Figure 1, thereby rendering the model more biologically realistic. First, inspired by [22], we retrained the network with the additional constraint that hidden unit activities be positive. This constraint can be implemented by simply changing the single neuron nonlinearity from hyperbolic tangent to a rectified linear unit (ReLU) (Figure 1C, black traces at left). Under these conditions, many units develop strikingly regular hexagonal grid maps similar to those of entorhinal grid cells (Figure 1C, lower), and the distribution of grid scores shifts to larger values (Figure 1E, and see also Supplementary Figure 2 for examples of model grid cells with different grid scores). Below, we present a mathematical theory that conceptually explains *why* networks with nonnegative firing rates develop hexagonal grid cells and explore the effect of nonnegativity constraints in a variety of network architectures. Second, we found that training with a small amount of weight decay on the recurrent weights encourages a path integration mechanism that continues to operate well beyond the walls of the training environment (Figure 1D, lower). Complementarily, we discovered a different training curriculum which achieves the same effect: rather than training in a single square environment, training consecutively in many environments with different geometries furnishes the trained network with the ability to generalize outside any one training environment (for details, see Methods 4.2). Since weight decay is easier to implement and has the additional desirable property of promoting simpler, low-rank connectivity, we study networks trained with weight decay in the remainder of this work. All subsequent analyses were performed on models with hexagonal grid maps and the ability to path integrate outside the training environment.

After demonstrating that training a simple RNN with nonnegative firing rates leads to the spontaneous emergence of hexagonal grid representations in its hidden units (Figure 1C, bottom), we tested whether this effect generalizes to different network architectures, and further explored the effects of the desired place cell output. We focused on three cases. First, we explored the feed-forward architecture of [22] which does not path-integrate, but does take place cell *inputs* and learns to generate grid cell outputs (Figure 2A, left). Second, we considered the complex path integrator of [31] which generates grid responses in a *disconnected* layer of neurons presynaptic to the desired place cell code and postsynaptic to a hidden recurrently connected network (Figure 2A, middle). This network also possesses complex nonlinear single neuron properties corresponding to long-short term memory (LSTM) cells [22]. Third, we considered our simple architecture which generates grid-like responses in a single hidden layer of recurrently connected neurons with simple sigmoidal or ReLU nonlinearities (Figure 2A, right) (see Methods 4.2 for details of all architectures and training).

**Figure 2:**
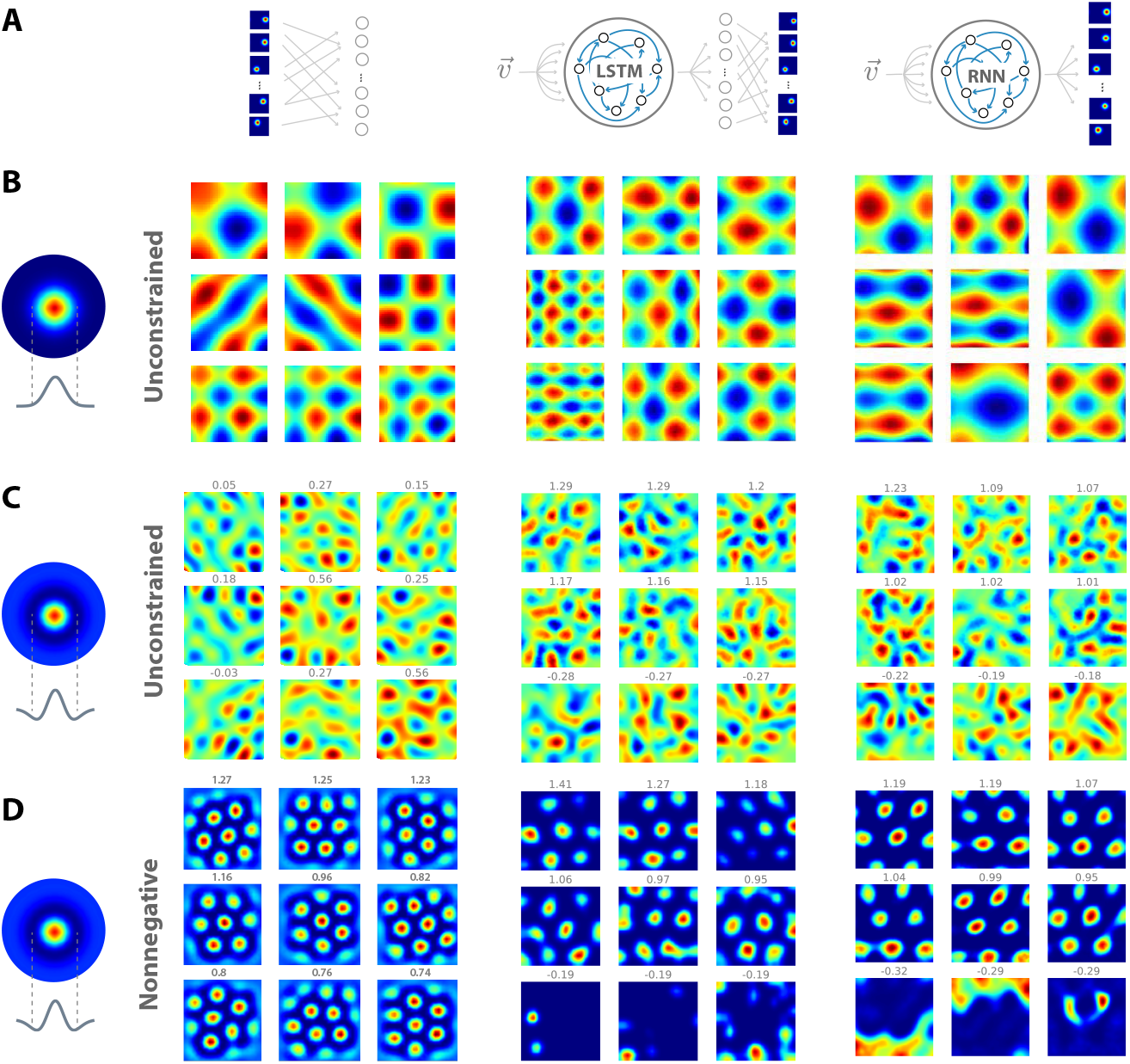
Neural networks trained on normative tasks develop grid-like firing fields. (A,B,C) From left to right, we train a single layer neural network, an LSTM, and an RNN on place cell outputs, reproducing the results of [22, 30, 31]. A) When the place cell receptive field (left) is broad, all networks learn square grids. B) When the place cell receptive field is a difference of gaussians, all networks learn amorphous, quasi-periodic patterns. C) When a nonnegativity constraint is imposed on hidden unit activations, all networks now learn regular hexagonal grids.

These three diverse architectures exhibited fairly universal properties of the learned hidden representations as a function of both the assumed place cell code, and the constraints on synapses in the feedforward model, or firing rates in the recurrent models. First, for wide place cell outputs without any surround inhibition, and with no constraints on synapses or firing rates, all three models learned *square* grid like responses (Figure 2B), similar to those found in [30]. If place cells have a surround inhibition, then all three models learn highly heterogeneous grid like responses (Figure 2C). In the feedforward model, this surround inhibition corresponds to place cell input firing rates suppressed below spontaneous rates, as in [22]. In the recurrent models, this surround corresponds to the layer of grid-like cells exciting the place cells with spatial RFs closest to the current position, and inhibiting the neighboring place cells. If in addition to the surround inhibition, we further constrain either synapses or firing rates to be non-negative, then all three models learn hexagonal firing fields (Figure 2D).

### 2.2 A theory for the emergence of diverse grid structures in trained neural circuits

The collection of training experiments in Figure 2 raises an intruiging question: why do these diverse neural architectures, across multiple tasks, all converge to near universal grid-like solutions, and what governs the lattice structure of this solution? We address this question by noting that all of the models described above contain within them a common sub-problem, which we call the position encoding problem. This problem involves selecting an optimal hidden representation that can generate place cell activity patterns through one layer of synaptic transformation. We develop our mathematical theory in Methods 4.4, and summarize its salient points here at a conceptual level. Overall, our theory allows us to understand the nature and structure of the resultant grid-like solutions in Figure 2, and their dependence on various modelling choices. Readers who are primarily interested in how hexagonal grid cell responses are mechanistically generated from the connectivity and dynamics of the learned circuit can safely skip this section.

A key quantity in our theory is the similarity structure of the output place cell representation, which we first define. Let *p_i_*(*x*) denote the activity of place cell *i* when the animal is at spatial location *x*. More precisely, *p_i_*(*x*) can be thought of as the input current to place cell i which is then rectified to generate place cell firing rates, so that the firing rates can only be positive while *p_i_*(*x*) can be negative. The similarity structure of the place cell code is then described by the spatial similarity matrix Σ_*xx*′_ = ∑_*i*_ *p_i_*(*x*)*p_i_*(*x*′). A positive (negative) matrix element Σ_*xx*′_ quantifies how similar (dissimilar) the place cell code is at two points *x* and *x*′ in space. Figure 3A shows a single row of the similarity matrix where *x* is the center of a 2D enclosure, and *x*′ varies over the enclosure, in the case where the place cell maps *p_i_*(*x*) have a center-surround structure as in Figure 2C,D. The similarity is high and positive for points *x*′ near the center, low and negative for points further away, and close to zero for points even further away.

**Figure 3:**
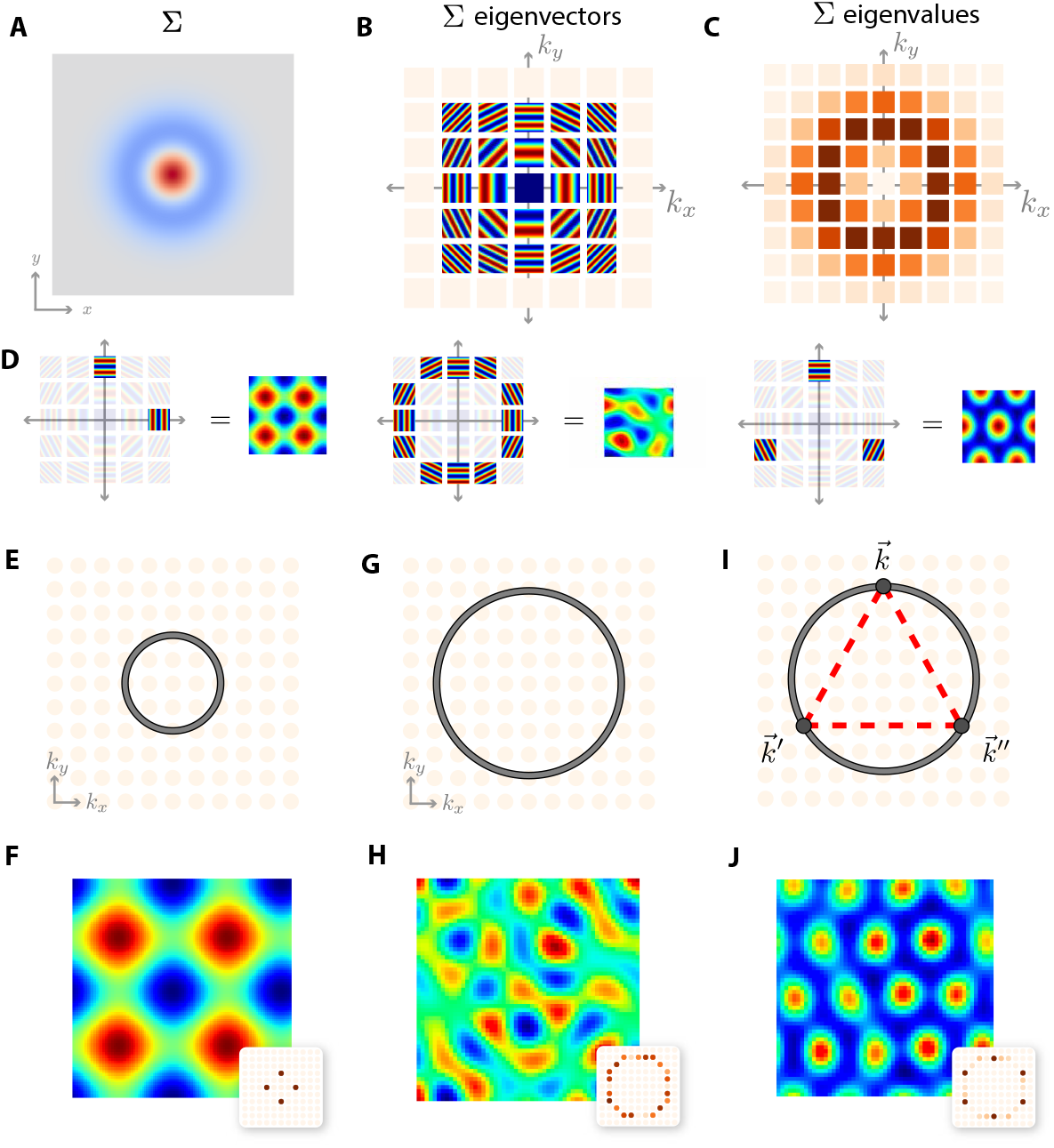
A theory for predicting the structure of learned spatial maps. (A) A central row of the place cell similarity matrix Σ, indicating how similar the desired place cell code at any point in a 2D enclosure is to the place cell code at the center, when place cells have a center surround structure as in Figure 2CD. Red, blue and grey indicate positive, negative, and zero similarity respectively. (B) Eigenvectors of Σ are well approximated by Fourier plane waves, and are shown arranged on a discrete lattice of integers (*k_x_, k_y_*) corresponding to the two oscillation frequencies of each plane wave along the two cardinal axes. (C) The eigenvalues of Σ corresponding to the eigenvectors in B. Darker red indicates larger eigenvalues. (D) Particular linear combinations of plane waves yield grid patterns. A square grid can be generated by two waves each oscillating along one of the cardinal axes (left). A heterogeneous grid can be generated by arbitrary linear combinations of plane waves with oscillations frequencies near an annulus of fixed radius (middle). A hexagonal grid pattern can be generated by a combination of plane waves whose oscillations frequencies form an equilateral triangle in the (*k_x_, k_y_*) plane, centered at the origin (right). (E) If the similarity structure in panel A is wide, the large eigenvalues of Σ will lie near an annulus of small radius, shown in grey, that only intersects a few plane modes oscillating primarily along the cardinal axes. (F) Simulations confirm that neural circuits with unconstrained firing rates will learn combinations of precisely these cardinal modes, generating square grids. (G) For a narrow similarity structure in panel A, the large eigenvalues of Σ will lie near an annulus of large radius, shown in grey. (H) Simulations confirm that neural circuits with unconstrained firing rates will then learn arbitrary linear combinations of plane waves with oscillations frequencies near this annulus, generating heterogeneous grids. (I) Adding a nonnegativity constraint forces neural circuits to learn combinations of plane waves whose oscillation frequencies sum to 0 as vectors in the (*k_x_, k_y_*) lattice. A simple such example is 3 plane waves whose oscillation frequencies lie at the vertices of a centered equilateral triangle (dashed red). (J) Simulations confirm that neural circuits with nonnegative firing rates will learn hexagonal grids, as predicted by theory in panels D (right) and H. In panels F,H,J, the learned grid cell representation is shown, with its Fourier power spectrum depicted as an inset.

According to our theory, the similarity structure Σ_*xx*′_ of the place cell code and the constraints on the hidden units determine what kind of grid cells emerge. Indeed, the similarity structure of inputs and outputs often determines the nature of learned representations in neural networks, from the development of ocular dominance columns [44] to the development of semantic categories [45, 46]. In our context, the eigenmodes and eigenvalues of Σ_*xx*′_ have a strong influence on the learned grid cell representations. The eigenmodes of Σ_*xx*′_ are spatial functions of 2D position and are well approximated by Fourier plane waves that oscillate in different frequencies and directions (Figure 3B, and Methods 4.4). The eigenmodes are indexed by two integers, *k_x_* and *k_y_*, indicating the spatial frequency of oscillation in each of the two cardinal spatial directions. Each such eigenmode has an associated nonnegative eigenvalue. These eigenvalues are shown in Figure 3C at the integer (*k_x_, k_y_*) lattice points associated with the corresponding eigenvectors in Figure 3B. The strength of these eigenvalues can be obtained by computing the power in each Fourier mode at spatial frequency (*k_x_, k_y_*) of the similarity function displayed Figure 3A, which corresponds to the central row of Σ_*xx*′_ (see Methods 4.4). Because this similarity function has a center-surround structure, the maximal Fourier power occurs near an annulus in the (*k_x_, k_y_*) lattice, and the narrower the similarity function in Figure 3A, the larger the radius of this annulus in Figure 3C.

The key significance behind the eigenvalues in Figure 3C, is that the learned hidden representations of neural networks will be preferentially composed of linear combinations of eigenmodes in Figure 3B with large associated eigenvalues (see Methods 4.4). Thus, while *any* function over the 2D enclosure can be written as *some* linear combination of Fourier eigenmodes, trained neural networks will only learn restricted linear combinations corresponding to a subset of these eigenmodes. Figure 3D indicates how, for example, square, heterogeneous, or hexagonal grid patterns can be constructed from appropriate combinations of Fourier modes. The key issue then is, how does the structure of the place cell code conspire with constraints on hidden representations to almost universally generate these three types of grid codes?

Our theory answers this question by elucidating three general scenarios that lead to these three qualitatively distinct types of codes. In the first case, if the place cell similarity structure in Figure 3A is wide relative to the size of the enclosure, then the maximal eigenvalues will occur near an annulus of small radius, as in Figure 3E. This annulus will intersect a small number of lattice points in the (*k_x_, k_y_*) plane corresponding to low frequency eigenmode oscillations aligned along the cardinal axes of the enclosure, and linear combinations of these oscillations along these cardinal directions would predict square grid cells as in Figure 3D, left. This prediction is confirmed in simulations of our position encoding problem in Figure 3F (see Methods 4.4). Indeed, square grid cells were previously found in trained path integrators [30]. On the other hand, if the similarity structure in Figure 3A is narrow, the maximal eigenvalues will occur near an annulus of large radius, which intersects many lattice points in the (*k_x_, k_y_*) plane, as in Figure 3G. With no further constraints, the hidden representation of neural circuits will learn arbitrary linear combinations of eigenmodes associated with the many lattice points on the large annulus, yielding relatively heterogenous patterns, as predicted in Figure 3D, middle. This prediction is confirmed in simulations in Figure 3H. Indeed [31] found highly heterogeneous grid-like representations, with a few cells having a high grid score, but the entire distribution of grid scores was indistinguishable from that obtained by grid-patterns obtained by low-pass filtering random noise, as demonstrated in Figure 1E.

If in addition to having a narrow place cell similarity structure (relative to the size of the enclosure), one also imposes a nonnegativity constraint on the firing rates of the hidden representation, we prove (see Methods 4.4) that in our position encoding problem, an additional constraint arises on which eigenmodes can be combined to construct hidden representations. As in the unconstrained case, hidden representations are learned by combining eigenmodes with large eigenvalues near the annulus in Figure 3I, but arbitrary linear combinations of all such eigenmodes are no longer allowed. Only linear combinations of eigenmodes whose weighted integer lattice points, as vectors in the (*k_x_, k_y_*) plane, sum to zero can be combined. The simplest such set is 3 lattice points forming an equilateral triangle centered at the origin, as shown in Figure 3I, and the combination of the associated eigenmodes predicts a hexagonal grid representation as in Figure 3D, right. This prediction is confirmed in simulations of our position encoding problem in Figure 3J.

Taken together, our theory provides a unifying conceptual explanation for when and why square, heterogeneous or hexagonal grids spontaneously emerge across three diverse architectures depicted in Figure 2A, and elucidates why prior work [30, 31] on training path integrator neural networks did not find truly hexagonal grid structure. We now move beyond the computational origin of grid cells to understanding the mechanistic origins of these cells in trained neural networks.

### 2.3 Two dimensional attractor dynamics underlies path integration in trained networks

Any neural circuit that is required to maintain a memory trace of position while the animal is standing still, with no velocity or other sensory inputs, must maintain a set of stable attractor patterns of neural activity. Many traditional hand designed neural network models for grid cells in two dimensions build in, *by design*, a 2*D* attractor manifold of stable activity patterns that has the structure of a torus [9–12, 20]. This raises a fundamental question: does a specifically toroidal manifold of attractor patterns occur naturally and generically across the space of neural networks trained to path integrate, or do such tori exist primarily only within the minds of theorists who design path integrators? We took advantage of having complete access to both the connectome and dynamics of our trained networks to search for and characterize neural activity patterns that are stable for long periods of time, using the methods of [47]. Remarkably, we found a large number of such attractor patterns that could be arranged, in a chosen 6 dimensional subspace, continuously along a 2*D* manifold with the shape of a torus (Figure 4A,B, see Methods 4.5 for details). Moreover, the attractor patterns were in correspondence with positions in physical space; smooth motions of the animal in physical space lead to smooth motions of the neural circuit dynamics along the attractor manifold on individual simulated trajectories (Figure 4B, dashed line), and the decoded position associated with each attractor pattern varied smoothly across the attractor manifold (Figure 4B, blue-green colormap). Moreover, we found that as the animal moves from one end of the enclosure to another, at least within this 6 dimensional subspace, the neural trajectory wraps multiple times around this 2D attractor manifold. Indeed, three different 2*D* projections of attractor manifold within the 6 dimensional space reveal this wrap around phenomenon. In each of the 2D projections, the neural trajectory wraps multiple times around a ring of the full torus as the animal moves along a direction that is either 0°, 60°, and 120° with respect to the horizontal axis of the enclosure (Figure 4D).

**Figure 4:**
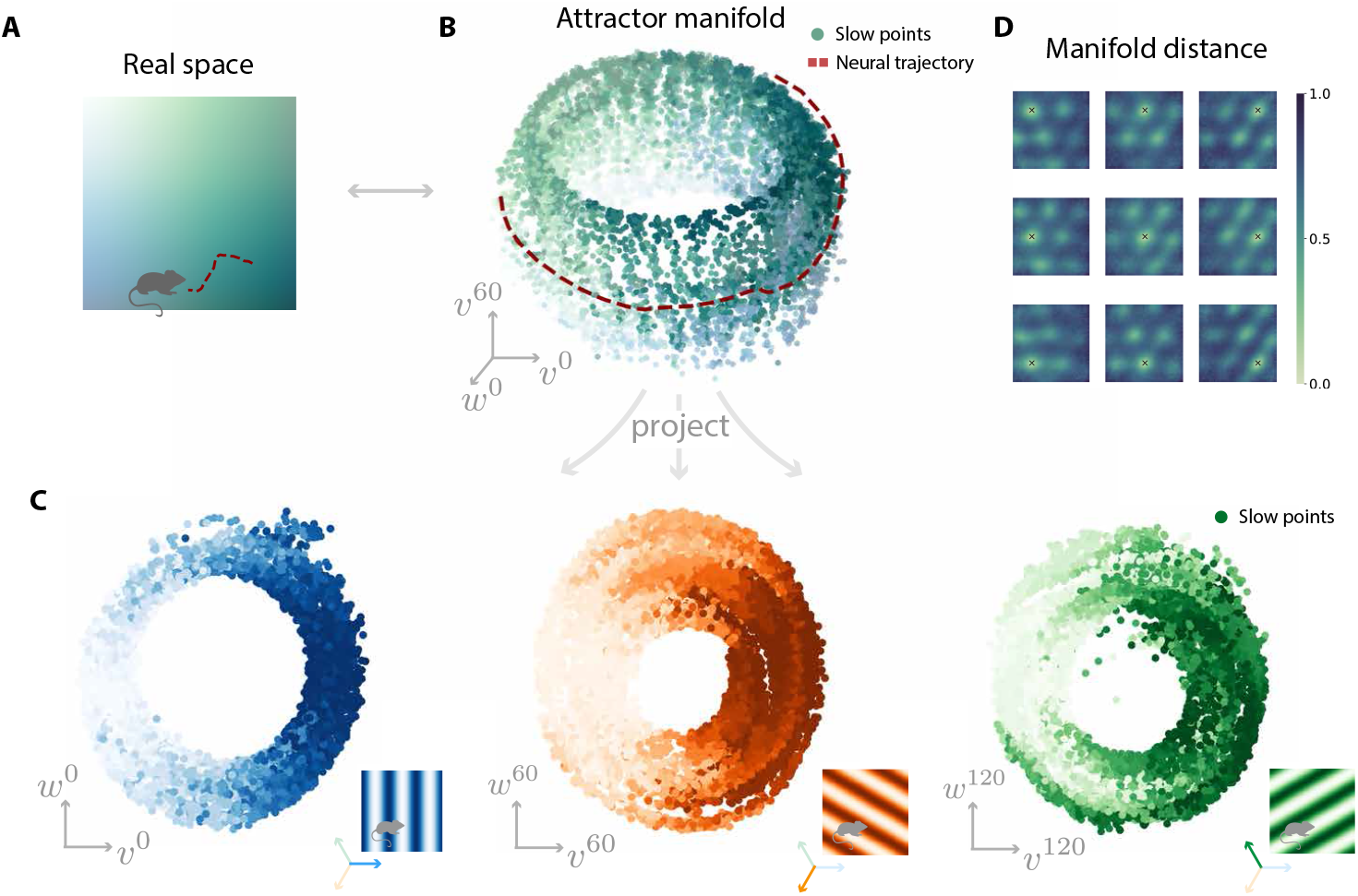
Emergent two dimensional attractor neural dynamics in trained path integrators. (A) An example trajectory (red dashed line) across a 2D environment. (B) A toroidal manifold of stable attractor patterns in neural activity space, visualized via dimensionality reduction, with each attractor pattern colored by the decoded position within the 2d environment in panel A. The neural trajectory corresponding to the animal trajectory in panel A is shown as a red dashed line. (C) Distance between the attractor pattern associated with a reference point in the 2d environment (marked with a black x) and the attractor patterns associated to all other points in the 2d environment. Different subpanels show different reference points. (D) Projecting attractor patterns onto three suitably chosen pairs of axes (see Methods 4.5) reveals three rings, representing position along the 0°, 60°, and 120° vectors in real space. Colors correspond to decoded position in the 2d environment (insets).

Thus, while we found the theorist’s Platonic ideal toroidal attractor manifold exists, at least within a 6 dimensional subspace of activity patterns, this analysis raises two fundamental questions. First, is the toroidal structure that we find simply an artifact of the subspace that we chose? We show this is not the case by repeating the same analysis that we did on the attractor patterns to derive Figure 4B,D, but this time we applied the analysis to random patterns obtained by low pass filtering spatial noise as in Figure 1D. The results, shown in (Supplementary Figure 1A,B), indicate a clear qualitative departure from a toroidal manifold structure. This control analysis indicates that our novel toroidal subspace finding method will not spuriously generate tori when they don’t exist, even in high dimensional datasets. Second, while we have shown this toroidal attractor manifold is not an artifact of our chosen 6 dimensional subspace, a natural question is, how does the 2D manifold structure behave outside of this subspace, and is it approximately toroidal in the full space? A key property of the toroidal structure is that as the animal moves along the 0°, 60°, and 120° directions in physical space, motion along the attractor manifold curls back to itself, to a very good approximation within the 6*D* space. We can also compute distances between different pairs of attractor patterns, not only in the 6 dimensional space, but also in the full space of all *N* = 4096 neurons, thereby not relying on dimensionality reduction. For each subpanel in Figure 4C, we pick a reference location in the spatial environment (indicated by a red ‘x’), and as the agent moves through the box, we build a heatmap showing the distance between the current neural activity pattern and the pattern active at the reference location. These distances indicate that as the animal moves in physical space preferentially along the 0°, 60°, and 120°, the attractor manifold does indeed approximately, but not fully, return to itself multiple times in the full space of all neurons. When the full distance computation analysis is applied to a subset of cells with high grid score, the manifold returns much more closely to itself (Supplementary Figure 1C), indicating that the partial departure from toroidal structure receives a contribution from neurons with heterogeneous neural activity patterns that simultaneously coexist with the neurons with more regular hexagonal grid patterns.

Thus overall, trained neural networks, while containing some of the structure of the perfect toroidal 2*D* attractor manifold that underlies hand-designed models, nevertheless also contain within them a much more general and varied 2*D* manifold structure that goes beyond the Platonic torus. This observation is intriguing because the principle of translation invariant connectivity, which leads to the perfect toroidal manifold, is a key principle theorists employed to design 2*D* attractor networks. Our exploration of a much larger class of networks that can path integrate reveals that the theorist’s perfect toroidal 2D attractor manifold solution is but one of many in a much larger space of neural circuit solutions to path integration. This larger space of network solutions includes many neurons with highly regular grid patterns that *simultaneously* coexist with many neurons with more heterogeneous patterns (Supplementary Figure 2), as is consistent with a recent statistical analysis of MEC firing patterns [48]. Interestingly, the hand design of path-integrator networks with a simultaneous coexistence of neurons with both highly structured and highly heterogeneous firing patterns has so far eluded the capabilities of theorists. But such networks are naturally found by neural network training, thereby validating such an approach as a powerful method for generating circuit level hypotheses for neural function with more biologically realistic levels of heterogeneity.

We next turn our attention to how circuit connectivity and dynamics conspire to both generate the 2D attractor manifold, and update position along this manifold as velocity signals enter the network. Below we review how traditional hand-designed models solve the two core problems of storage and updating of position, and reveal how trained networks solve these same problems. By comparing them, we find that trained networks find more general classes of solutions than those found in theorists’ hand-designed models, while still retaining some of their core computational elements, albeit in modified forms. We begin by considering path integration in one dimension, as the results are easier to visualize and understand, before moving on to two dimensions.

### 2.4 Stable storage of angular information in one dimension

To elucidate the circuit mechanisms underlying path-integration in trained networks, we began by training the neural architecture in Figure 1B to instead integrate a 1-dimensional periodic variable, corresponding for example to head direction, with angular head velocity as input. In this section we ask how this trained circuit connectivity maintains a stable memory trace of head direction, and in the next section we ask how this trace is updated when nonzero angular velocity inputs arrive.

Traditional head-direction path integrator models [13–17] consist of a line (or more precisely a ring) of neurons that can maintain a one parameter family of stable, localized bump attractor patterns that occupy different positions on the neural line (Figure 5A (see Methods 4.6 for simulation details). All of these bump attractors are stabilized by a translationally invariant synaptic connectivity matrix with local short-range excitation and long-range inhibition (Figure 5B). To more clearly see salient features in this connectivity, we can also look at the connectivity in the Fourier basis of periodic waves on the ring (Figure 5C) as opposed to the neuron basis (Figure 5B) (see Methods 4.8). Intuitively, the connectivity matrix element *J_ij_* in the neuron basis tells us how strongly an input to neuron *j* elicits activity in neuron *i*. Similarly, the connectivity matrix element *J_ωω′_* in the Fourier basis tells us how strongly a periodic wave of stimulation on the neural line at frequency *ω′* elicits a wave like response at frequency *ω*. Because the connectivity of hand-designed models is translation invariant, the connectivity in the Fourier basis will be diagonal, meaning that input patterns at a given frequency *ω* only influence output patterns at that same frequency at any appreciable strength. Furthermore, in the simplest model capable of maintaining a bump of stable activity patterns, namely the ring attractor [49], only two periodic patterns exist in the connectivity: the sine and cosine functions of lowest frequency. Thus in this model, as shown in Figure 5C, the connectivity is exceedingly simple: it only has two nonzero elements reflecting the fact that the sine and cosine patterns of lowest frequency are the only ones that self-sustain themselves.

**Figure 5:**
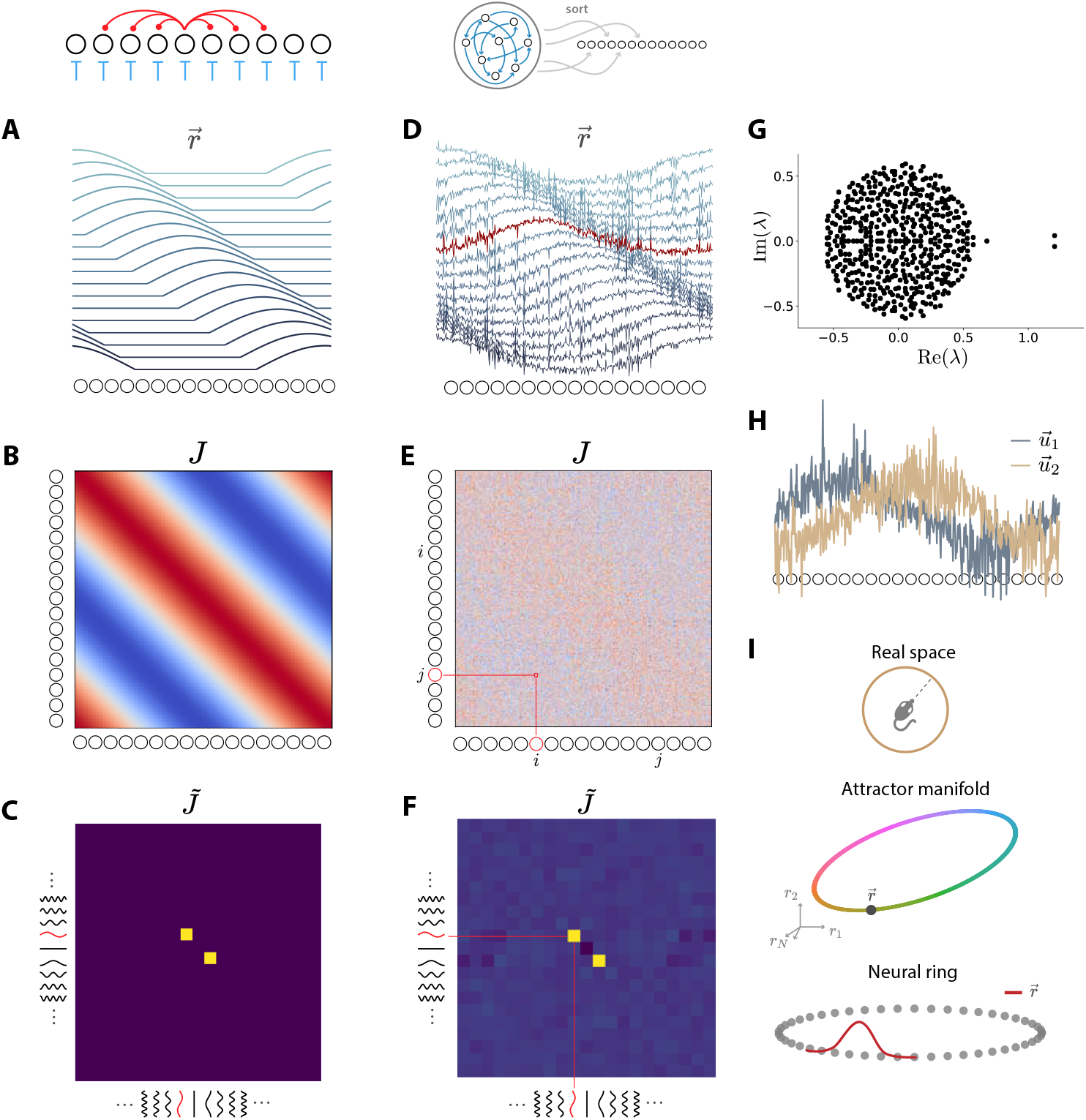
Mechanisms of information storage in one dimension in hand-designed and trained neural networks. (A) Hand-designed networks employ a 1D ring, or line of neurons, with local excitation and long range inhibition. This connectivity results in a set of stable localized bump activity patterns (blue curves) on the 1D neural ring (see Methods 4.6 for simulation details). (B) The synaptic connectivity matrix between pairs of neurons sorted according to position on the neural ring; red (blue) indicates excitation (inhibition). (C) The same synaptic connectivity matrix as panel B, but now the connections are between all pairs of Fourier modes on the neural ring, with the lowest frequency modes at the central rows and columns. The only two nonzero elements correspond to the sine and cosine patterns of lowest frequency providing self-excitation to themselves. (D) We can disentangle neurons in a trained recurrent network to order the neurons along a neural ring by sorting the neurons from left to right according to the preferred head direction. We then find the stable activity patterns of the network, and sort them from top to bottom according to the peak of their excitation on the neural ring. (E) The connectivity matrix of the trained network when neurons are sorted according to their position on the extracted neural ring. (F) The same connectivity as in panel E but now between Fourier modes on the extracted neural ring. (G) Eigenvalues of the trained connectivity matrix. (H) The eigenvectors associated with the two outlier eigenvalues in panel G roughly correspond to sine and cosine functions on a neural ring that can be derived from the connectivity alone (see Methods 4.7). (I) Visualizing information storage in three different spaces. (Upper) The animal at some heading in real space. (Middle) The current neural activity pattern corresponds to a point on a ring manifold of stable attractor patterns in neural activity space. (Lower) This pattern is a localized bump pattern on the neural ring.

We next asked, what elements of this highly structured ring attractor solution depicted in Figure 5A-C are present, if at all, in trained path integrators. First, in a trained network, the neurons do not possess any natural ordering. We thus first sorted the neurons via their preferred head direction by measuring their tuning curves, yielding a neural line. We also found and sorted the stable activity patterns, revealing a surprisingly well behaved set of localized activity bumps (Figure 5D, see Methods 4.7 for details). Encouraged by this match in activity, we asked whether the network’s connectivity matrix, again with neurons sorted according to the neural line derived from tuning curves, resembles the center-surround connectivity of the idealized model (Figure 5B). Despite faint banding, there is little obvious structure present in the network’s recurrent interactions (Figure 5E), at least in the neuron basis. The relative lack of structure serves as an instructive lesson for connectomics efforts. For example, any noise in the estimation of synaptic connectivity strengths may render such structure completely invisible to analysis efforts that operate only within the single neuron basis. However, despite the lack of obvious structure in the trained network’s weight matrix in the neuron basis (Figure 5E), in the Fourier wave basis it strikingly recapitulates the simple structure of the hand-designed ring attractor (Figure 5F).

Our ability to reveal the simple structure of the connectome Figure 5F relied on our ability to sort the neurons with knowledge of all stable activity patterns as in Figure 5D. Instead of using neural activity to organize the connectome, can we also extract this simple connectivity structure from the knowledge of the connectome *alone* and use it to organize activity? To investigate this possibility, we computed the eigenspectrum of the connectome (Figure 5G), revealing two large eigenvalues. We used the corresponding eigenvectors to sort the neurons on a connectome derived neural line (as opposed to the activity derived neural line in Figure 5D). When we plotted the associated eigenvectors, along this neural line, we again found approximate sine and cosine patterns along the neural line (Figure 5H). These sine and cosine patterns hiding within the dominant modes of the connectome, ultimately contribute to the ability of the network to maintain a family of stable bump patterns in Figure 5D, which can be thought of as different linear combinations of the sine and cosine patterns. Thus, in this case at least, knowledge of the connectome alone can be used to approximately understand and organize activity patterns, as long as one focuses on the dominant modes of the connectome, and not the individual synaptic strengths. A natural question is how did training develop this dominant mode structure? Strikingly, we find that training amplified a nascent mode structure present in the randomly initialized connectivity to arrive at this final connectivity (See Supplementary Figure 3 and Methods 4.8).

In summary, we have thus seen that the trained network stores an estimate of current head direction as an activity bump along an emergent neural line (Figure 5D), and such activity bumps are stablized by dominant connectivity modes (Figure 5G). Such a family of bump patterns can be thought of as a ring attractor manifold of stable activity patterns in firing rate space, as shown schematically in Figure 5I. Each head direction in real space (inset) is represented by a location on the attractor ring in activity space (top), and in turn, each point on the ring corresponds to a specific bump pattern along the neural line (bottom).

### 2.5 Velocity based updating of angular information in one dimension

We next asked how the trained RNN updates its stored head direction as it receives angular velocity inputs. Hand-designed path integrator models accomplish this updating through two subpopulations of neurons with offset outgoing connections, biased either rightward or leftward. Velocity signals indicating that the animal is turning right (left) preferentially target the neural population with rightward (leftward) biased projections, which cause increased activation on the right (left) of the current activity bump, thereby shifting this bump right (left) (Figure 6A; see Methods 4.6 for simulation details). This process can be understood more precisely by linearizing the dynamics while the network is at a stable bump pattern and a velocity input *v* is given to the network. This linearization leads to a sequence of steps. First the velocity input *v* drives each neuron *j* in the attractor network through a feedforward synaptic strength *M_j_*. This feedforward drive is multiplied by the gain 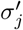 of neuron *j*, where 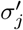 is the slope of the neuron’s input-output curve while the network is in a stable bump pattern. Then this feedforward drive propagates through the recurrent connectivity *J_ij_* to yield a net excitation or inhibition to every neuron *i* in the network given by 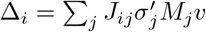 (see full derivation in Methods 4.9). This process is depicted graphically in Figure 6B for a hand-designed network where the rows of the recurrent connectivity *J_ij_* are sorted by the sign of the feedforward synapses *M_j_*, and the columns of the connectivity are sorted by the preferred head direction. There are clearly two populations of cells marked by whether a cell is excited (inhibited) by the velocity input with a positive (negative) feedforward synapse *M_j_*. Moreover, cells in the population with a positive (negative) *M_j_* are precisely the cells that have a rightward (leftward) biased set of outgoing connectivity *J_ij_*. This clear conspiracy in the connectome between a cell’s feedforward drive from velocity, and biased outgoing connections to the rest of the neurons, leads to a result in which the net drive Δ_*i*_ will excite (inhibit) the rightward (leftward) edge of the current activity bump when the velocity is positive, thereby moving the bump right. Similarly, the bump will move left when the velocity is negative. (Though connectivity alone is sufficient to understand the model in this case, the drive term Δ_*i*_ tells us that more generally, one has to consider the nonlinearity *σ* as well. Indeed, as we show in Supplementary Figure/Video 4, changing the nonlinearity from ReLU to tanh surprisingly causes the network to integrate in the opposite direction - that is, stimulating cells with rightward-biased connectivity causes the population activity pattern to move left, and vice-versa - an interesting and potentially important cautionary tale for efforts aimed at understanding circuit function on the basis of connectivity alone.)

**Figure 6:**
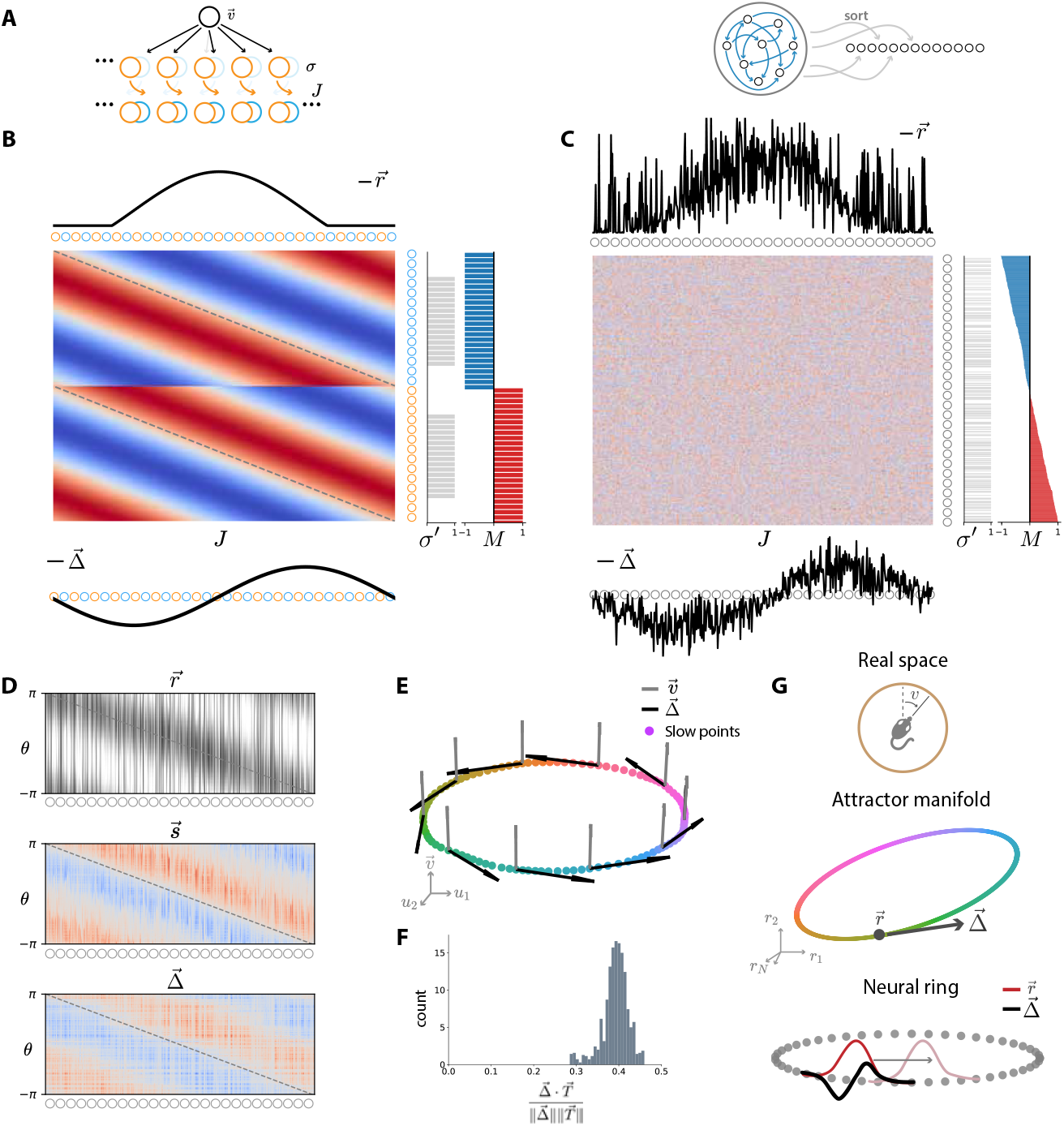
Mechanisms of one dimensional path integration in hand-designed and trained neural networks. (A) Schematic of a connectivity rule for updating position in many hand-designed models. An incoming velocity input (black cell marked 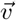) excites (inhibits) an orange (blue) population of cells that have biased rightward (leftward) outgoing recurrent connectivity (see Methods 4.6 for simulation details). (B) Illustration of velocity integration in a hand-designed network. The current activity pattern is stable bump on the neural ring (top). When the velocity cell is activated, through a set of feed-forward synapses *M_j_* it sends excitation (red bars) to right-shifting cells and inhibition (blue bars) to left-shifting cells. This excitation and inhibition passes through the single neuron gains 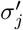 (grey bars). This modulated excitation/inhibition passes through the recurrent connectivity matrix *J_ij_*, shown with the rows sorted by strength of the feedforward synapses *M_j_* and the columns sorted by position on the neural ring. The combined result is a pattern Δ_i_ of excitation (inhibition) (black curve bottom) provided to the leading (lagging) edge of the current activity bump, moving it to the right for a positive velocity input *v*. (C) We show the exact same quantities that were plotted for the hand-designed network, but now for the trained network, where the neural ring is extracted by sorting neurons according to their preferred head directions. Notably, there is a single continuum of positive (negative) feedforward synaptic strengths shown by the red (blue) bars on the right, and on a neuron by neuron basis, these feedforward synapses are not strongly related to the recurrent connectivity (lack of discerinble structure in *J_ij_*). However, the total pattern of excitation and inhibition Δ_*i*_ in the black bottom curve is similar to that of the hand designed network in panel B. (D) (Top) Stable activity patterns along the neural ring of the trained network at all head directions. (Middle) The spatial derivative of these activity pattern along the neural ring, with respect to head direction. (Bottom) The pattern of excitation and inhibition Δ_*i*_. The qualitative match between this latter pattern and the spatial derivative indicates that the excitation/inhibition patterns will cause roughly uniform translation along the attractor manifold for all head directions. (E) The set of stable activity patterns in the trained network visualized in a 3 dimensional space spanned by the top 2 principal components of neural activity and the velocity input vector *M_j_*. The excitation/inhibition pattern Δ_*i*_ points tangentially along the ring attractor manifold at all locations. (F) A histogram of the cosine angle between the tangent vector to the ring of stable patterns 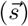 and the pattern of velocity driven excitation/inhibition (Δ_*i*_) indicates it is approximately constant along the ring, further demonstrating velocity inputs push the neural state along the ring at a constant rate. (G) Visualizing position updates in 3 different spaces. (Top) The animal turns in real physical space. (Middle) This turn yields a net pattern of excitation/ inhibition Δ_*i*_ that is tangent to the ring of stable neural attractor patterns. (Bottom) The current stable attractor pattern *r_i_* is a bump on the neural ring (red curve) while the current velocity driven excitation/inhibition pattern Δ_*i*_ is a approximately a derivative of this bump on the neural ring (red curve). Adding this latter pattern in moves the bump.

Does the trained RNN update its representation of head direction using the same simple shifting mechanism? We sorted the connectome of the trained network in Figure 6C using exactly the same rules we used to sort the connectome of the hand-designed network in Figure 6B; the rows are sorted by the strength of the feedforward connectivity *M_j_*, from most negative to most positive and the columns are sorted by the preferred head direction of each neuron. We first note that the feedforward synaptic strength *M_j_* does not divide the neurons into two clear populations; rather this strength varies along a continuum. Second, despite the sorting of neurons, the recurrent connectome *J_ij_* bears no clear relation to the feedforward connectome *M_j_* on a neuron by neuron basis for any individual neuron *j*, as it does in Figure 6B. However, once one sums over the collective effects of all neurons *j* to obtain the net drive to each neuron *i*, again given by 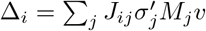, we see a clear pattern in Δ_*i*_ in which the one side of the bump is excited and the other side is inhibited, causing a net coherent motion of the current activity bump.

In Figure 6D-F we confirm that the net drive Δ_*i*_ appropriately shifts the bump not only at one location (as in Figure 6C) but all locations. The network updating operation can then be viewed geometrically as motion along a ring of stable activity patterns in firing rate space, with the net drive Δ_*i*_ acting as a vector that is tangent to the ring at all points along the ring (Figure 6G, middle). In summary, the trained network mimics the overall design of the hand-designed network, at the key functional level of net overall excitation Δ_*i*_, but it does not implement this function in a clearly discernable way on a neuron by neuron basis in the connectome. Instead, the trained RNN takes advantage of many small non-systematic outgoing projection biases and combines them into a coherent drive to move the current bump in the appropriate direction. Thus individual synaptic components do not have a clear functional meaning, but the collective dynamics does. If evolution does indeed find more generic network solutions to path-integration as in Figure 6C (compared to less generic solutions as in Figure 6B), then the observed emergence of dynamic function through the collective effects of many synapses working together again poses an instructive lesson for connectomics efforts that might seek to find function in the presence or absence of individual synapses.

### 2.6 Stable storage of positional information in two dimensions

We now turn to the problem of storing position in two dimensions. Traditional models accomplish this by arranging neurons on a two dimensional neural sheet, with a center-surround connectivity over this sheet (Figure 7A), yielding a set of stable hexagonal firing patterns on the neural sheet (Figure 7B; simulation details in Methods 4.6). Because every single cell has outgoing projections onto its neighbors in the neural sheet organized in a center-surround fashion, the averaged outgoing connectivity of each cell to its neighbors has a clear center surround structure (Figure 7C, left). This connectivity structure implies that the top eigenmodes of the connectivity, when displayed as patterns on the neural sheet, are approximately Fourier modes. Thus if we plot the connectivity from each Fourier mode to itself in the 2D Fourier basis on the neural sheet (see Methods 4.8 for details), we see that a small number of Fourier mode patterns on the neural sheet self-excite themselves (Figure 7C, right). Moreover, these top 6 eigenmodes of the connectivity are shown in Figure 7D as patterns on the neural sheet, and they clearly represent low frequency sine and cosine waves along three different directions. Thus a 2D family of stable patterns is maintained fundamentally through the self excitation of low frequency Fourier modes along the 2D neural sheet.

**Figure 7:**
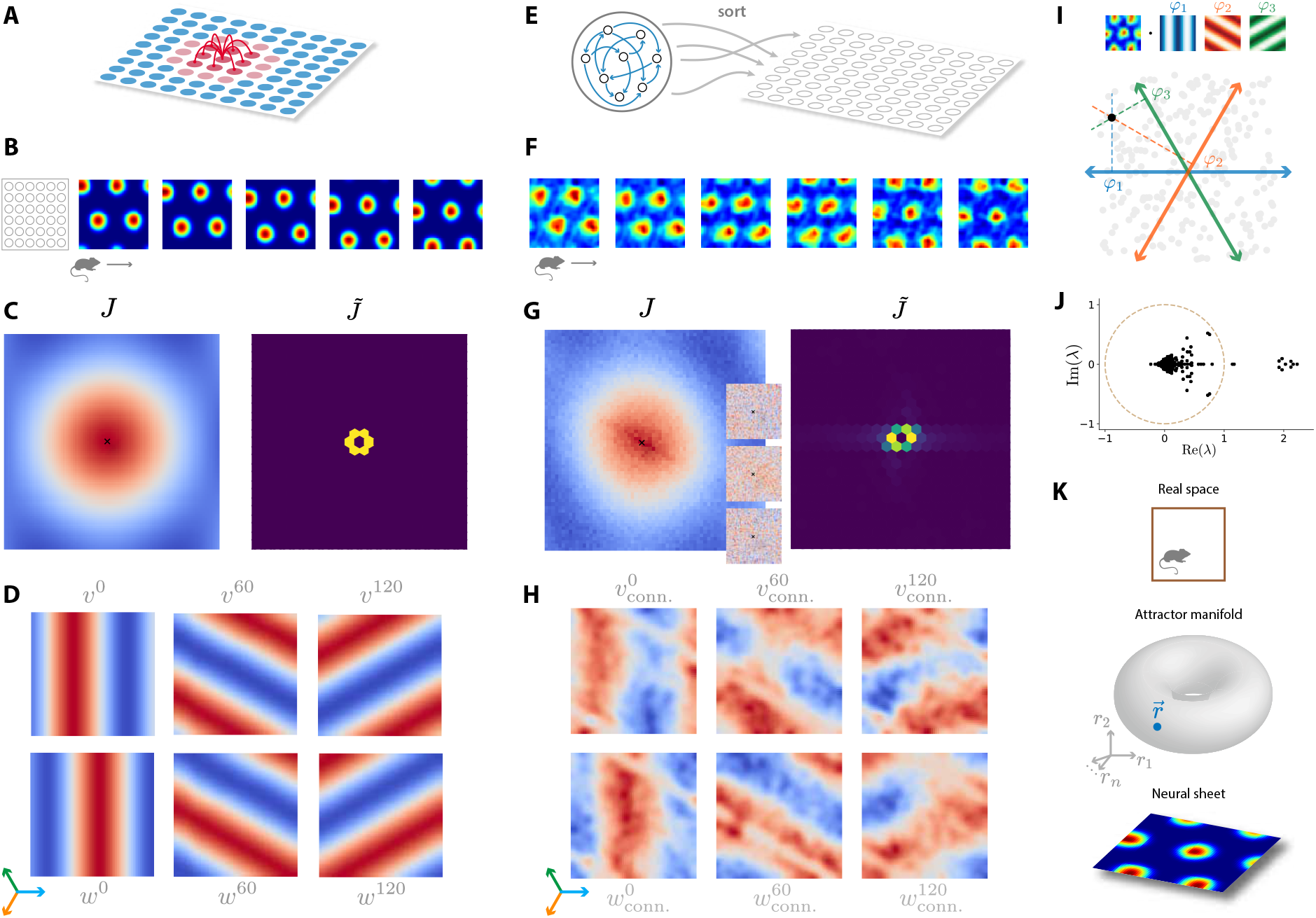
Mechanisms of information storage in two dimensions in hand-designed and trained neural networks. (A) Hand-designed networks employ a 2D sheet of neurons, with local excitation and long range inhibition to yield stable activity patterns (simulation details in Methods 4.6). (B) Five different stable activity patterns on the neural sheet are shown when the animal is at 5 successive positions in physical space. (C) The average outgoing connectivity profile (red (blue) indicates excitation (inhibition)) over all neurons to all other neurons at a given displacement on the 2D neural sheet (left), and the degree of self-excitation provided by Fourier modes on the 2D neural sheet to themselves through the connectivity. Fourier modes on the neural sheet are indexed by 2 discrete frequency variables, just as Fourier modes in physical space are indexed by two discrete frequency variables in Figure 3B,C. The connectivity is such that a small number of low frequency Fourier modes excite themselves. (D) The top 6 eigenmodes of the connectivity correspond to low frequency Fourier modes on the 2D neural sheet. (E) A schematic depiction of our analysis method to disentangle neurons in a trained recurrent network to order the neurons along a 2D neural sheet to reveal mechanisms underlying path integration in trained networks. (F-H) Plots of same quantities as in panels B-D, except now the quantities are computed for the trained network using the 2D neural sheet extracted by our analysis (see Methods 4.7 for sort details). The additional 3 insets in panel G left show three example outgoing connectivity profiles for three neurons, revealing no discernible structure. Structure in the connectivity only appears after averaging outgoing connectivity profiles across neurons (G, left) or in the basis of Fourier modes (G, right), both as functions on our extracted 2D neural sheet. (I) A schematic of our method to position neurons on a 2D neural sheet. For each cell we compute the phase of the Fourier transform along 3 different axes. For hexagonal patterns, these phases obey a linear relation enabling us to explain them using only two variables, which then become coordinates on a neural sheet. For more heterogenous cells, we find the best coordinates we can to explain the 3 phases, thereby placing every neuron (grey dot) at some point on a 2D neural sheet. This analysis method of organizing neurons in a low dimensional space is crucial to revealing the emergent structure of trained neural networks in panels F,G,H. (J) Eigenvalues of the trained connectivity, again indicating positive feedback for a small number of eigenmodes. (K) Visualizing the storage of position in three different spaces. (Upper) The animal at a location in physical space; (Middle) The current neural activity pattern corresponds to a point on a 2-dimensional toroidal manifold in neural activity space; (Lower) The current neural activity pattern can be visualized as a pattern of hexagonal bumps on a 2D neural sheet.

Thus hand-designed networks for storing position in two dimensions build in a significant amount of structure, as depicted in Figure 7A-D. On the other hand, it is *a priori* unclear how the connectivity of trained neural networks enable them two maintain a family of stable hexagonal firing patterns of many individual cells across physical space. A key impediment to understanding this lies in the absence of any organizing principle to sort the neurons onto a putative neural sheet, and to understand the connectivity as a function of the relative position of pairs of neurons on the neural sheet. In one dimension we were able to easily sort neurons by their preferred head direction to obtain a neural ring (Figure 5C). But this simple strategy does not generalize to two dimensions because single neuron firing rate maps are now multi-modal and also many are significantly heterogeneous. We were thus faced with the challenge: is there a procedure for physically organizing a population of hexagonal grid cells that will reveal structure in their connectivity and activity patterns? And under this organization, do the simple mechanisms used in continuous attractor models (Fig. 7A-D) qualitatively match those learned by the trained network? To address these issues, we developed a new, noise-tolerant sorting procedure tailored to hexagonal maps (Fig. 7I). Briefly, for each rate map, we measured 3 spatial phases designed to locate the neuron’s hexagonal grid along the 0°, 60°, and 120° axes. We then arranged the neurons onto a 2D neural sheet such that neurons with similar spatial phases were physically close (see Methods 4.7 for full details). This method enables us to use single neuron activity patterns to untangle the trained neural network and organize the neurons on a 2D sheet (Fig. 7E).

After sorting the neurons, when we plot stable activity patterns of the trained network as patterns over the sorted 2D neural sheet, we obtain, remarkably, hexagonal firing patterns over the emergent neural sheet (Figure 7F). We note that these hexagonal firing patterns are distinct from, but related to, the firing fields of individual grid cells across physical space, as shown for example in (Fig. 2D). The latter involve the average firing rate of *single* cells across *all* of physical space, while the former involve the activity of *all* neurons across a neural sheet, while the animal is at a *single* point in physical space, analogous to a single frame in a Calcium imaging experiment [50].

Now that we have used neural activity to organize neurons on a 2D neural sheet, we can examine the neural connectivity. The insets in Figure 7G show three examples of the outgoing connectivity strength of a single neuron at the center, to every other neuron as a function of the relative displacement of each recipient neuron on the neural sheet. These insets show very little structure. However, when this displaced connectivity pattern is averaged over all neurons, we see a clear center-surround structure on average (Figure 7G, left). Moreover, when we examine how strongly each Fourier mode on the neural sheet maps to itself through the connectivity, we find that a small number of Fourier modes excite themselves (Figure 7G, right). We also computed the top eigenmodes of the connectivity matrix. While each eigenmode was not a clear Fourier mode on the neural sheet, we found linear combinations of the top 10 eigenmodes (see Methods 4.10 for details) that, when plotted as functions on the neural sheet, exhibited clear Fourier wave like structure, resembling sine and cosine plane waves oriented at 0°, 60°, and 120° degrees (Figure 7H). Moreover, the eigenvalues of the connectivity are shown in Figure 7J), revealing a small number of strong eigenmodes in the connectivity. Thus, analogous to the 1D network in Figure 5I, as the animal traverses the 2-dimensional environment, the pattern of neural activity can be thought of as a point on a stable manifold in neural activity space, or as a hexagonal grid of activity clusters on a neural sheet (Figure 7K).

In summary, remarkably, the trained neural network finds a solution to the problem of storage of 2D position in a manner that is quite similar, at a collective level, to the hand-designed neural network, as seen by the qualitative similarity of (Fig. 7A-D) and (Fig. 7E-H). This conclusion was not *a priori* obvious, and required analysis methods that: (1) employ activity to organize neurons on a neural sheet, and (2) find combinations of connectivity eigenmodes that behave simply as functions on the neural sheet. Notably, individual synaptic strengths have a much less clear meaning (Figure 7G, insets). Instead, our ability to understand the essential principles underlying memory storage in this network required the combined analysis of both activity patterns and the connectome.

### 2.7 Velocity based updating of positional information in two dimensions

Hand-designed path integrator networks update their stored position using a mechanism (Figure 8A; simulation details in Methods 4.6) that is analogous to the one used in 1D (Figure 6A). Now, instead of two populations of neurons with biased left/right outgoing projections, there are four populations of neurons with biased outgoing recurrent projections in the north, south, east and west directions on the 2D neural sheet. Linearizing the dynamics as we did in the 1D case, leads to a net pattern of excitation or inhibition provided to the network given by 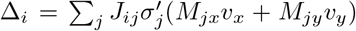, where *v_x_* and *v_y_* are the firing rate of *x* and *y* selective velocity input neurons respectively (full derivation in Methods 4.9). The feedforward synapses *M_jx_* and *M_jy_* to each neuron *j* determine the preferred velocity of that neuron *j*. To update position correctly, hand-designed networks are organized such that each neuron’s preferred velocity matches its biased pattern of outgoing connectivity. For example, a neuron *j* with feedforward synapses yielding north, east, west, or south velocity selectivity also has biased outgoing connections in the same direction on the neural sheet. This structure yields a situation in which animal motion in any direction in physical space moves the bump on the neural sheet in a corresponding direction, through a net excitation/inhibition pattern Δ_*i*_ that leads to excitation (inhibition) of neurons at the leading (lagging) edge of the current activity bump along the correct axis on the neural sheet (Figure 8B). As shown below, the dynamics of these hand-designed networks can be thought of as motion along a toroidal manifold of stable activity patterns that is pushed along this manifold in a velocity dependent manner (Figure 8C), resulting in translation of a hexagonal pattern on the 2D neural sheet (Figure 8D).

**Figure 8:**
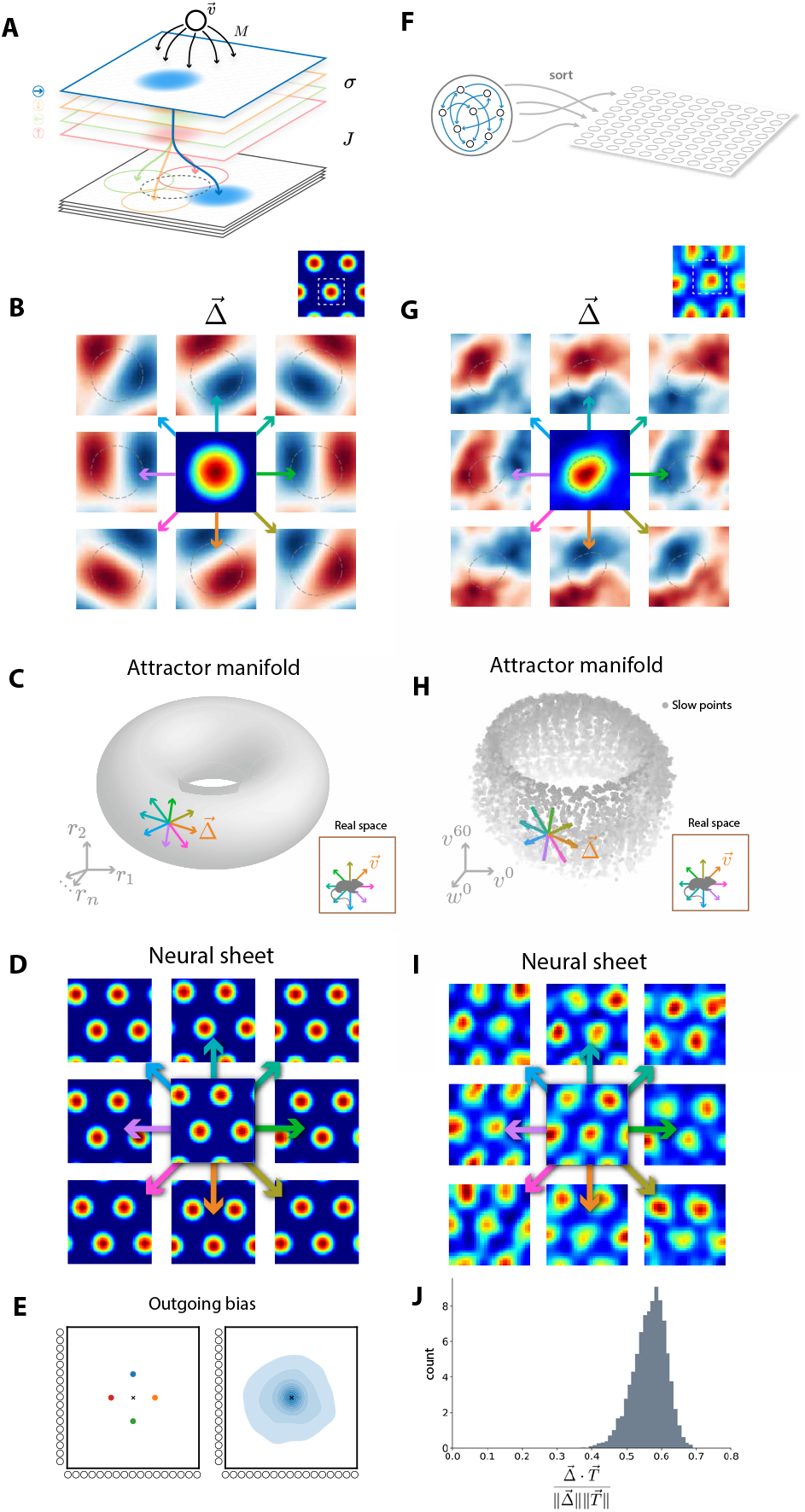
Mechanisms of two-dimensional path integration in hand-designed and trained neural networks. (A) Schematic of connectivity rule for updating position in many hand-designed models (full details in Methods 4.6). Incoming velocity inputs (black cell) marked 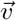 with feedforward synapses *M_j_* (black arrows), excite a population of eastward-projecting neurons (blue). The active eastward-projecting neurons (blue circle, top) excite population of neurons (blue circle, bottom) to the east of the current bump of activity (dashed grey circle, bottom), through eastward-offset recurrent connectivity (blue arrow, *J*), shifting the bump of activity to the east. Had the velocity inputs instead targeted south, west, or northward projecting neurons (orange, green, red), the bump would have instead shifted south, west, or north. (B) The pattern of excitation (red) and inhibition (blue) Δ_*i*_ provided to each neuron on the neural sheet when the animal is moving in eight directions in physical space for a hand-designed network. The central pattern is a small section of activity on the neural sheet before the motion, corresponding to the white dashed box in a larger cross section of the neural sheet (upper right). (C) While the animal moves in eight different directions in physical space (bottom right), the corresponding excitation/inhibition pattern Δ_*i*_ can be thought of as a tangent vector pointing along the 2D toroidal manifold of stable activity patterns, starting from the current pattern. (D) Motion of the hexagonal pattern on the 2D neural sheet for each of the 8 motions of the animal in physical space. (E) A histogram of outgoing biases in connectivity for each neuron, as a function of the relative displacement on the neural sheet, both for hand-designed (left) and trained (right) networks (see Methods 4.11). (F) A schematic of our novel disentangling procedure to take a trained network and organize neurons along a 2D neural sheet (see Figure 7I and Methods 4.7). (G,H,I) Same as (B,C,D) except all quantities are plotted as a function of position on a 2D neural sheet extracted from a trained network, as opposed to one designed by hand. (J) Histogram of the cosine angle between the pattern of excitation inhibition Δ_*i*_ generated by a velocity input, and the correct direction of motion along the attractor manifold, across all points on the manifold. For appopriate path-integration, this cosine angle need not be 1, but should be roughly constant across the manifold, as it is.

Thus hand-designed networks build in a significant amount of structure in the feedforward and recurrent connectivity. How do trained networks then update position? By leveraging our ability to untangle trained networks and sort their neurons onto a 2D neural sheet using activity patterns alone (Figure 7E,I), we examined the histogram of biases in the outgoing connectivity of each neuron to all other neurons on the neural sheet. For a hand designed network, this histogram would yield four populations of biases in outgoing projections (Figure 8E, left). However, in trained networks, we found the histogram of biases was much more random and unimodal, centered around zero bias (Figure 8E, right; details in Methods 4.11). Nevertheless, after sorting the units onto a neural sheet (Figure 8F), when we computed the main quantity that determines function, namely the network pattern of excitation and inhibition delivered across the neural sheet, again given by 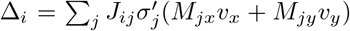, we found this overall pattern, obtainable only by summing over all neurons *j* in the network, behaved as in the hand-designed network, providing the correct excitation (inhibition) pattern to neurons at the leading (lagging) edge of the current activity bump along the correct axis on the neural sheet (Figure 8G). This yields a geometric picture of the operation of the trained network in which the stable activity patterns form a toroidal manifold in the same 6 dimensional subspace we found in Figure 4B,D, and the pattern of excitation/inhibition Δ_*i*_, when reduced to this space, pushes activity tangent to the manifold along a direction determined by the velocity in physical space (Figure 8H). This push moves the hexagonal pattern on the neural sheet in the correct direction (Figure 8I), and indeed the cosine of the angle between the push given by Δ_*i*_ and the tangent direction of correct motion along the manifold stays roughly constant along the entire manifold of stable activity patterns (Figure 8J).

Thus overall, at the level of the collective dynamics, the trained network has hiding within it the same computational principles of a 2D neural sheet and correct pushes by velocity along this sheet (compare Figure 8B,G). However, unlike the hand-designed network, there is much more heterogeneity at the level of individual synapses (compare Figure 8E, left and right). Again, in more generic solutions found by neural network training, the principles underlying network function emerge at the collective level, but remain murky at the level of individual components (synapses and neurons).

## 3 Discussion

In summary, we have demonstrated that neural networks trained to path integrate, under simple biological constraints, can robustly generate hexagonal firing fields and can generalize outside of their training environment. Moreover, we develop a general unified theory for *why* grid cells with different lattice structures can spontaneously emerge in diverse normative as well as mechanistic models. Finally, we develop novel algorithmic methods to extract, from the seemingly highly unstructured connectomes of such trained networks, a conceptual understanding of how such networks both path integrate as well as mechanistically generate hexagonal grid cell responses. These results address and raise a host of interesting issues, both for the specifics of understanding entorhinal function, and about general methods of arriving at a conceptual understanding of circuit function in neuroscience from large scale datasets and models.

### 3.1 From function to structure in MEC

The maxim of Francis Crick, “If you want to understand function, study structure,” reflects a common approach to gaining knowledge in biology by first examining the structure inherent in a biological system, and then inferring from that structure, its function. In the case of MEC we have long had two disparate ideas: first, MEC grid cell firing patterns have a striking hexagonal structure, and second we have long hypothesized that MEC plays a role in path integration [5, 51]. However, the precise relation between these two ideas has remained unclear. In this work, we have worked backwards from function to structure. Indeed we have forged a link between the computational function of path-integration and the structure of hexagonal grid cell patterns by demonstrating that neural circuits with simple constraints on firing rates naturally generate hexagonal grid cells when forced to path-integrate. Moreover, we have developed a general mathematical theory explaining *why* this happens. By showing that hexagonal grid patterns arise as a structural consequence of the function of path-integration, we provide further evidence that this function may have guided the early evolution and development of MEC circuitry. Of course, MEC could have further evolved to implement even more general functions, as we discuss below.

### 3.2 On order and randomness in network function

A striking difference between traditional hand designed models of MEC function [9–12] and neurobiological reality likes in the crystalline precision of grid cells typically realized in the former, and the substantial heterogeneity of neural firing patterns, including those of grid cells, found in the latter [48]. To date, no theoretical models have been able to simultaneously capture the coexistence both structured hexagonal grid cells, and heterogenously firing grid cells in the same interconnected neural population. Through training, our models achieve exactly this, thereby opening the possibility of exploring a much larger space of circuit hypotheses for entorhinal function that more faithfully captures its underlying heterogeneity. In particular these models raise intriguing theoretical questions about how structured and random firing patterns can coexist in the same interconnected network, and open up the possibility of connecting theory and experiment in a much more detailed manner than traditional hand designed models that do not capture heterogeneity.

### 3.3 Whither lies function: from connectomes and activity maps to conceptual understanding

Substantial experimental effort has gone into generating connectomes [52–55] and brain activity maps [56], but it is unclear how we should analyze such large scale data, and at what level of organization circuit principles will become manifest [41]. On the one hand, clues to circuit function may already appear at the level of individual neurons and synapses, and may be clear from direct inspection of the connectome, as in potentially highly specialized systems like the fly head motion system [57–59]. But in more generic situations, it is unclear what type of circuit solution evolution itself finds to any given computational problem, and whether more indirect relations between connectomes, brain activity maps and circuit function may exist that are not apparent at the level of individual neurons and synapses. The process of training neural networks to solve any given computational task then provides an excellent basis for exploring the space of circuit solutions. However, the development of analytic methods to bridge the gulf between trained connectomes and activity maps, and conceptual understanding then becomes essential [34, 41, 47]. We have developed such tools for understanding how trained path integrators work, and we find in many cases that both the connectome and brain activity maps can yield insight into circuit function, though not at the level of individual neurons and synapses. However, conclusions drawn from connectomes alone could be incorrect, and in the settings analyzed, brain activity maps provide more accurate views on circuit function (e.g. Supplementary Figure 4). In general, we hope our attempts to understand how trained networks function in the case of path integration will inspire similar efforts in other tasks, thereby generating new hypotheses for circuit operation in diverse settings.

### 3.4 Future directions

In general, our work raises several possibilities for interesting future directions. First, it would be interesting to incorporate landmarks, in addition to velocity inputs, as a secondary source of information about position. The fusion of landmark and velocity inputs has been previously studied in hand-designed models, which have successfully accounted for the deformation of grid cells in irregular environments [12] and the remapping of grid cells in virtual reality environments [21]. However, such models, with their crystalline grid-cell structure, cannot make predictions for what heterogeneous cells would do under the same experimental manipulations. Intriguingly, non-grid spatial cells have been shown to remap more readily to environmental manipulations than grid cells [60]. Second, recent work has shown that changes in rewarded location can affect grid cell firing properties [61, 62]. Training neural networks to forage in response to changed rewarded locations could yield new, general hypotheses about interactions between reward, spatial location, and grid cell and heterogeneous cell firing patterns.

Perhaps more ambitiously, fundamental new ideas may be required to obtain an understanding of the functional role of MEC grid cells in more general settings beyond that of path integration, for example in general episodic memory, abstract reasoning, planning, and imagination [29, 63, 64]. Indeed, ancient mechanisms for integrating velocity to arrive at position, in path integration, may have been co-opted by evolution to integrate individual deductive steps to arrive at final inferences, in general processes of reasoning and imagination. To obtain mechanistic hypotheses as to how neural circuits can accomplish such remarkable feats, the proximal path may lie in training neural circuits on complex tasks, developing analytic tools to understand their function, and extracting predictions that can be verified at emergent levels of circuit organization, beyond individual neurons and synapses, in large scale connectomes and brain activity maps. We hope our work along these lines in the simple setting of path integration inspires similar ways forward in more complex settings.

## Acknowledgements

We would like to thank Shaul Druckmann for helpful discussions. We also thank the following for financial support: the Stanford Graduate Fellowship for B.S., the Stanford Neurosciences Graduate program for G.M., an Urbanek fellowship for S.O., the Office of Naval Research N00141812690, Simons Foundation SCGB 542987SPI, NIMH MH106475, and the James S McDonnell Foundation for L.M.G., and the Simons and James S McDonnell foundations, and an NSF CAREER award for S.G.

## 4 STAR Methods

### 4.1 Notation

*n_g_* number of hidden recurrent neurons
*n_p_* number of output place cells
*n_x_* number of spatial arena locations
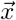 position in 2d physical space (arena)
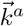 unit vectors in *a* = 0°, 60°, 120° directions.
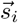 position of neuron *i* on neural sheet
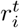 activity of neuron *i* at time *t*
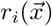 activity of neuron *i* at arena location 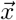
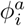 measured phase of neuron *i*’s spatial rate map in 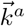 direction
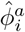 ideal phase of neuron *i*’s spatial rate map in 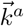 direction
*β_i_* outgoing connectivity bias of neuron *i*

### 4.2 RNN training

#### Path integration task

The task and training protocol shown in 1 were replicated from [31]. Place cell receptive field centers 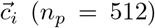 were distributed randomly over a (2.2*m* × 2.2*m*) environment. The response of the *i^th^* place cell was simulated using a difference of gaussians tuning curve 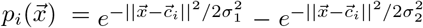 where *x* is the current location of the agent, and *σ*_1_ and *σ*_2_ represent the width of the center and surround, respectively. Agent trajectories were generated using the rat motion model described in [65]. Velocity signals from the simulated trajectory were given as input, and the network was expected to produce the simulated place cell activities as output.

#### Network architecture

The trained RNN studied throughout this work (cf. sections 2.3–2.7) is a “vanilla” RNN with the familiar discrete-time dynamics used in conventional ring attractor networks:

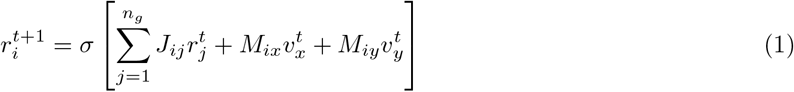

where *r^t^* is the population activity at time *t, J* is the (*n_g_, n_g_*) recurrent connectivity matrix, 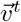 is the 2-dimensional velocity input at time *t, M* is the (*n_g_*, 2) matrix of velocity input weights, and *σ* is a pointwise nonlinearity (either tanh or relu). Predicted place cell outputs 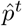 are read out linearly by a (*n_p_, n_g_*) matrix of weights *W*:

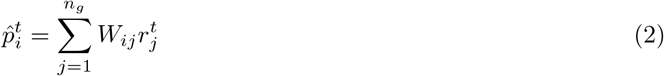

Rather than tuning the weights *J, M, W* by hand, we allowed them to be trained by gradient descent on the objective of reconstructing the true place cell outputs *p^t^* as accurately as possible. The loss function we used for training was a cross-entropy loss. Task, architecture and training hyperparameters are collected in the table below:

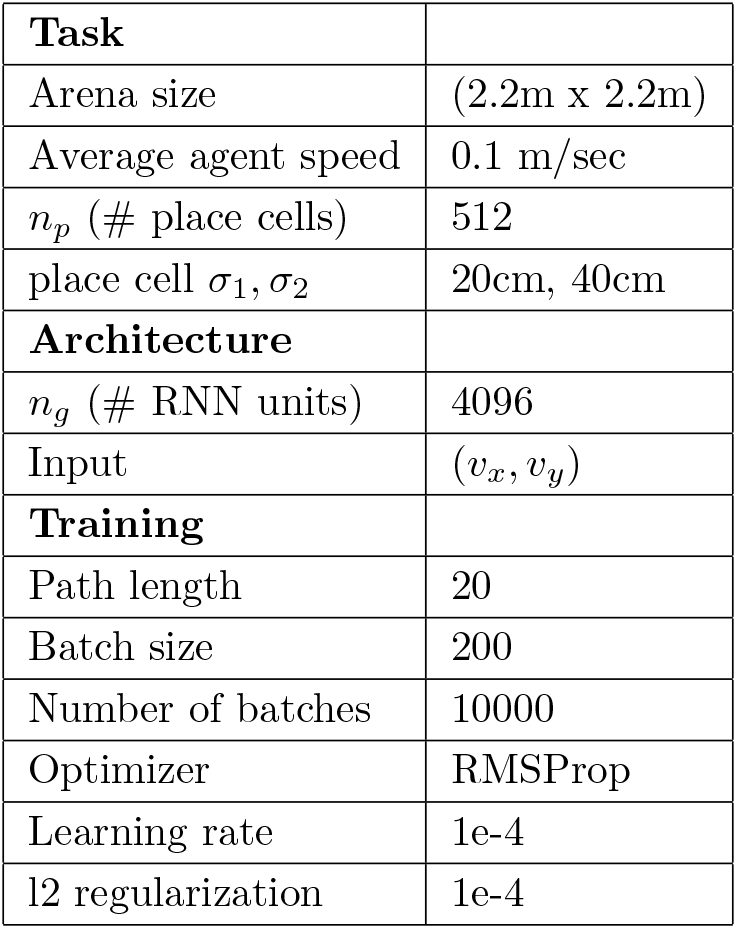

#### 1d RNN

Head direction (HD) networks (sections 2.4, 2.5 and Figures 5,6) were trained on a simplified 1D circular version of the spatial navigation task (described below). We simulated 32 head direction cells with preferred HD evenly spaced over 360 degrees. HD cell responses as a function of true head direction were computed as *h_i_*(*θ*) ∝ exp(*κ* cos(*θ* – *θ_i_*)) with *κ* = 2 and *θ_i_* is the preferred head direction of HD cell *i*. Agent heading trajectories were generated by first sampling turning velocity at each step from a Gaussian distribution with *μ* = 0 and *σ* = 0.8 radians. Velocity trajectories were then smoothed over time with a gaussian filter with *σ* = 1 step, and integrated to give head direction as a function of time. The network was given the trajectory velocities as input, and was expected to produce the responses of the 32 simulated HD cells as output. We used the same “vanilla RNN architecture as for the 2D integration task, but with *n_g_* = 512 hidden neurons rather than 4096.

#### 1-layer neural network

The 1-layer neural network shown in Figure 2B solves the problem of optimally reconstructing place cell activities *P* from *k* encoding maps *G* using readout weights *W*:

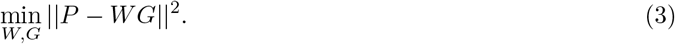

Without any additional constraints, this just amounts to low rank matrix factorization. By the Eckart-Young-Mirsky theorem, the optimal low rank *WG* has the same left and right singular vectors as *P*, and its singular values are obtained from *P*’s singular values by keeping the top *k* values and zeroing all others. This gives a particular *W* and *G*. All other optimal solutions can be obtained via the transformation *W* → *WM*, and *G* → *M*^−1^*G* for some invertible matrix *M*. The optimal maps *G* shown Figure 2B, left were obtained following the above by computing the singular value decomposition of *P* and extracting the top *k* = 9 left singular vectors.

With positivity constraints on *W*, and *G*, the problem becomes nonnegative matrix factorization. We used a standard toolbox for nonnegative matrix factorization (scikit-learn) for *k* = 9 maps. The resulting maps *G* are shown in Figure 2D, left.

#### LSTM path integrator network

The task and training protocol were identical to that of the RNN described above. The model architecture shown in Figure 2 was replicated from [31], consisting of x- and y-velocity inputs to an LSTM with 512 hidden units, followed by a linear layer of 128 units (which the authors called the “g-layer”), followed by a final readout to the estimated place cell activities. We trained both with and without an additional output to a population of head direction cells, as the authors in [31] did, and obtained similar results. We report results for the network that was trained to predict only place cell outputs. A summary of the task, architectural, and training parameters is given below:

| Task |  |
| --- | --- |
| Arena size | (2.2m x 2.2m) |
| Average agent speed | 0.1 m/sec |
| $n_p$ (# place cells) | 512 |
| place cell $\sigma_1, \sigma_2$ | 20cm, 40cm |
| Architecture |  |
| $n_g$ (# LSTM units) | 512 |
| # g-layer units | 128 |
| Input | $(v_x, v_y)$ |
| Training |  |
| Path length | 50 |
| Batch size | 200 |
| Number of batches | 10000 |
| Optimizer | RMSProp |
| Learning rate | 1e-4 |

#### Biological constraints

We imposed the following biologically inspired constraints on the RNN analyzed throughout the paper (sections 2.3–2.7) Nonnegativity: In order to achieve regular hexagonal grids, as our theory predicts, we imposed a nonnegativity constraint on the activities of the recurrent units in the RNN. We imposed this constraint by simply swapping the tanh nonlinearity for a relu (cf. Figure 1C), though we found that softer versions of this constraint, such as adding a penalty on negative firing rates to the loss function, also achieved the same effect.

Weight decay: We found that a small penalty 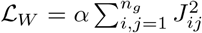 on the magnitudes of the recurrent weights encouraged a representation that generalizes beyond the boundaries of the training environment (Figure 1F). We simply added this penalty to the loss function for *α* = 10^−5^.

Multi-task learning: Similarly, we found that training in environments of many different geometries sequentially encouraged a representation that generalizes beyond the boundaries of any one training environment (Figure 1F). This learning curriculum was generated as follows: (1) construct a rectangular box with random width and height, (2) tile the box randomly with simulated place cells, (3) train in this environment for 1000 steps of gradient descent, (4) repeat.

### 4.3 Grid score

Grid score was evaluated as in [31]. A spatial ratemap was computed for each neuron by binning the agent’s position into 2cm× 2cm bins, and computing the average firing rate within each bin. Grid score was evaluated by rotating a circular sample of the spatial autocorrelogram of this ratemap in steps of 30°, and computing the correlation between the rotated map and the original. The grid score was defined as the minimum difference between the correlation at the expected peaks, (60°, 120°), and the correlation at the expected troughs (30°, 90°, 150°). The distribution of grid scores for all hidden units in the sigmoidal network with unconstrained firing rates, the null model, and the ReLU network with nonnegative firing rates are shown in Figure 1E.

### 4.4 Pattern formation theory predicts structure of learned representations

All of the trained models discussed in this work face essentially the same encoding problem of choosing *n_g_* encoding maps that generate *n_p_* place cell spatial maps through a single layer of synaptic weights (graphically depicted in Figure 1). We formalize this objective mathematically as follows:

Define the following matrices: 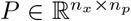 contains the responses of the *n_p_* place cells at all *n_x_* spatial locations; 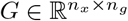 contains the responses of the *n_g_* encoding cells at all *n_x_* spatial locations; 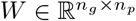 contains the readout weights from encoding cells to place cells. The goal is then to minimize

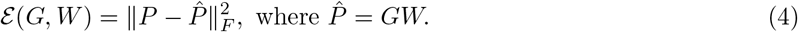

Here we consider an L2 penalty on encoding errors because it simplifies the analysis, but we observe similar results for numerical simulations with a softmax cross-entropy loss (see Figure 3).

Because we would like to understand the dominant patterns learned by the hidden neurons *G*, we make two simplifications to the above objective. First, we replace *W* by its optimal value for fixed *G*:

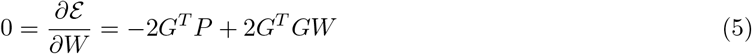

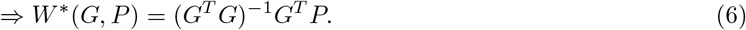

Observing that the objective 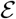 in (4) is invariant to any invertible transformation *Z* of the form *G* → *GZ, W* → *Z*^−1^*W*, we can simplify our objective by choosing *Z* so that *G*’s columns are orthonormal. Plugging (6) into (4) (and multiplying by a factor of 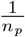 for convenience), we obtain the following constrained optimization problem for *G*:

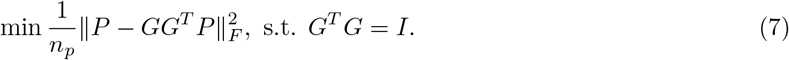

This constrained optimization problem can be solved by considering the Lagrangian

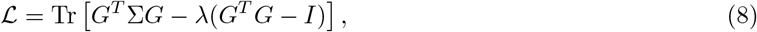

where 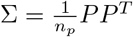 is the *n_x_* × *n_x_* place cell similarity matrix (shown in Figure 3A).

To build intuition, we consider the optimal pattern learned by a *single* hidden neuron. Replacing *G* with *g*, an *n_x_* × 1 vector of activations of a single hidden neuron at all points in space, we obtain the single-neuron Lagrangian

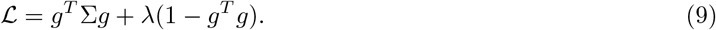

This is the simplest version of the position encoding objective. If we model the training of our neural network as performing gradient ascent on this objective, then the learning dynamics take the following form:

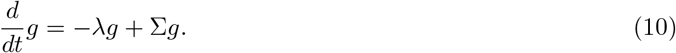

This is a **pattern forming dynamics**. As gradient ascent proceeds, the firing fields at two locations *g_x_, g_x′_* will mutually excite (inhibit) one another if the place cell similarity Σ_*xx*′_ at the two locations is positive (negative), and over time *g* will develop stable patterns across space. Solving these dynamics subject to the normalization constraint *g^T^ g* = 1, we find that the stable fixed point corresponds to the top eigenmode of Σ.

The top eigenmodes of Σ then take a very simple form. Assuming the place cell receptive fields uniformly cover space, then in the limit of many place cells, their similarity structure Σ is translation invariant: 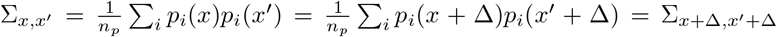. Without the boundaries, or with periodic boundary conditions on the box, this translation invariance would imply that Σ’s eigenvectors are exactly Fourier modes across space. However, even with the boundaries, Σ_*x,x′*_ has a Toeplitz structure and its eigenmodes are still well approximated by Fourier plane waves across space (Figure 3B). To compute the eigenvalue λ_*k*_ associated to the *k^th^* Fourier mode *f^k^*, let *p*(*x*) be the place cell tuning curve over space, and Δ_*i*_ be the receptive field center of the *i^th^* place cell. Then

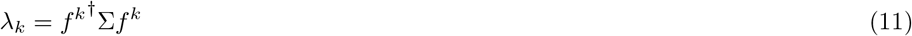

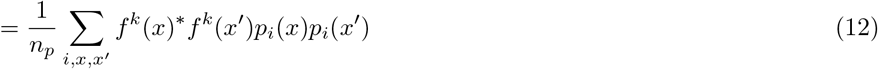

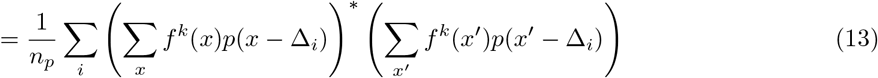

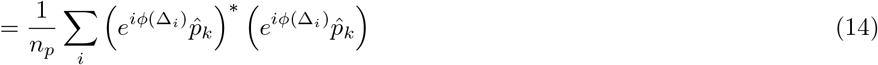

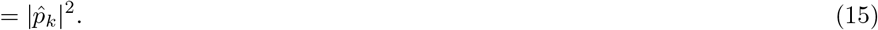

In essence, the eigenvalue associated to the *k^th^* Fourier mode is just the power of that Fourier mode in the place cell tuning curve, so that the optimal pattern *g* will be the Fourier mode with maximum power.

If the place cells are distributed isotropically across space, and their tuning curve is circularly symmetric then Σ_*Rx,Rx′*_ = Σ_*x,x′*_ for any rotation matrix *R*, and consequently all rotations of the optimal Fourier mode will also be optimal:

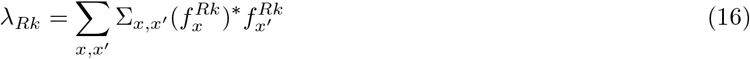

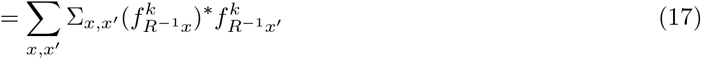

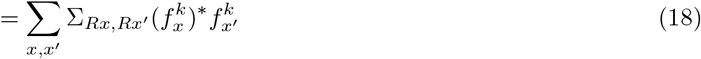

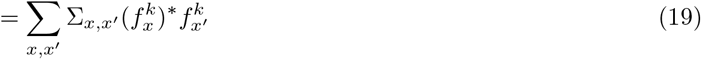

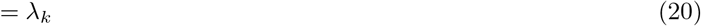

Thus, the top eigenspace of Σ is degenerate and consists of all Fourier modes whose wavevector *k* lies on a ring centered around the origin in Fourier space (Figure 3C). In other words, the optimal map *g* is any linear combination of plane waves of optimal wavelength 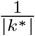, which can combine to form square, or hexagonal or even amorphous grid maps (Figure 3D). As we show below, this multiplicity of solutions is a special feature due to the lack of constraints. Once a nonlinear constraint such as non-negativity is added, the optimization favors a single type of map corresponding to hexagonal grid cells.

#### A nonnegativity constraint favors hexagonal grids

We have seen empirically that a nonnegativity constraint tends to produce hexagonal grids (Figure 2D). To understand this effect, we add a softened nonnegativity constraint to our objective function as follows

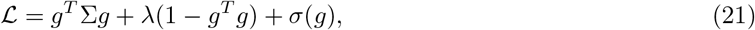

where *σ*(*g*) penalizes negative activities in the map *g*. It will be convenient to write *g_x_* as 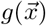, treating *g* as a scalar field defined for all points in space. Our objective then takes the form

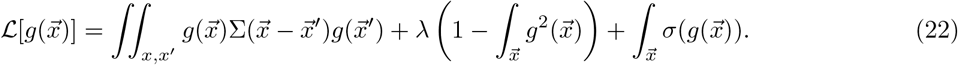

We can approximate the negativity penalty by Taylor expanding about 0: *σ*(*g*) ≈ *σ*_0_ + *σ*_1*g*_ + *σ*_2*g*_^2^ + *σ*_3*g*_^3^. Our Lagrangian then has a straightforward form in Fourier space

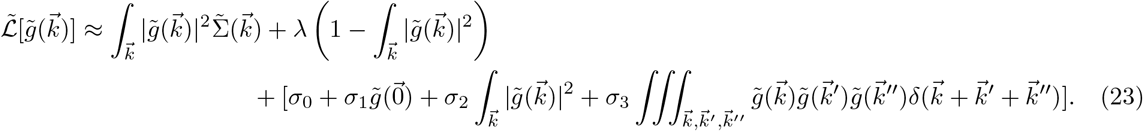

*σ*_0_, *σ*_1_, and *σ*_2_ will not qualitatively change the structure of the solutions: *σ*_0_ simply shifts the optimal value of 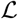, but not its argmax; *σ*_1_ controls the amount of the constant mode in the maps, and does not affect their qualitative shape; and *σ*_2_ can be absorbed into 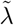 [66]. Critically, however, the cubic term *σ*_3_ introduces an interaction between wavevector triplets 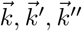 whenever the three sum to zero (3I).

In the limit of weak *σ*_3_, the maps will be affected in two separate ways. First, weak *σ*_3_ will pull the maps slightly outside of the linear span of the optimal plane-waves, or eigenmodes of Σ of largest eigenvalue. As *σ*_3_ → 0, this effect shrinks and effectively disappears, so that we can assume the optimal maps are still constrained to be linear combinations of plane waves, with wave-vectors on the same ring in Fourier space. The second, stronger effect is due to the fact that no matter how small *σ*_3_ is made, it will break 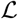’s symmetry, effectively forcing it to choose one solution from the set of previously degenerate optima. Therefore, in the limit of small *σ*_3_, we can determine the optimal maps by considering which wavevector mixture on the ring of radius *k*\* maximizes the nonlinear term

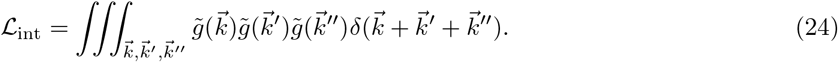

Subject to the normalization constraint 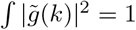, this term is maximized when 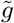 puts all weight on a single wavevector triplet which sums to zero: 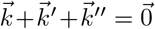. The only such combination on the ring of radius |*k*\*| is an equilateral triangle, so that the optimal solutions are 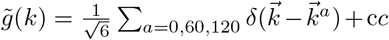.^1^ (Figure 3I). Therefore, rather than arbitrary linear combinations of plane waves, the optimal solutions consist of three plane waves with equal amplitude and wavevectors that lie on an equilateral triangle.

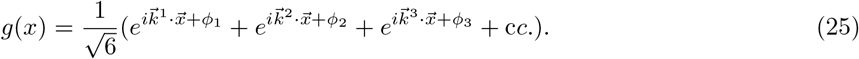

The interaction 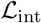 is maximized when *ϕ*_1_ + *ϕ*_2_ + *ϕ*_3_ = 0, in which case the three plane waves interfere to form a regular hexagonal lattice (Figure 3J).

#### Hexagonal grids and *g* → −*g* symmetry breaking

We see from the above argument that the rectification nonlinearity is but one of a large class of nonlinearities which will favor hexagonal grids. A generic nonlinearity with a non-trivial cubic term in its Taylor expansion will break the *g* → −*g* symmetry, and introduce a three-body interaction which picks out hexagonal lattices. While nonnegativity is a specific nonlinearity motivated by biological considerations, a broad class of nonlinearities will achieve the same effect [67].

#### Numerical simulation of pattern-forming systems

As we show above, the problem of optimally encoding place cell activities yields the following Lagrangian:

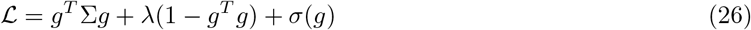

where Σ = *P^T^ P* is a (*n_x_* × *n_x_*) matrix encoding the similarity of the place cell activities at any given pair of spatial locations, and *σ* captures the nonnegativity constraint by penalizing negative activities.

Place cell tuning curves are modelled as a difference of gaussians as above in the path integration task:

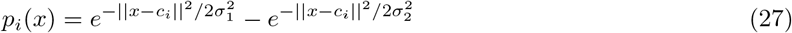

where *c_i_* is the location of the *i^th^* place cell’s receptive field center, and *x* is the current location of the agent. We sampled these tuning curves (*n_P_* = 512) on a grid of spatial locations (*n_x_* × *n_x_* = 100 × 100) to obtain the matrix of place cell responses, *P*, and computed Σ = *P^T^ P* (for difference of gaussian tuning curves, it is possible to obtain an approximate analytical expression for Σ; see appendix for details).

In the unconstrained case *σ* = 0, the optimum can be obtained directly by sampling from the top eigenspace of Σ. This is shown in Figure 3B,C. For other choices of *σ*, we numerically optimized 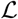 via gradient descent: 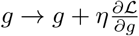. Optimization was run until approximate convergence. Figure 3J shows one such map for *σ* = relu(*x*).

### 4.5 Attractor manifold analysis

For the analysis of Figure 4, fixed points of the network dynamics are defined as population activity patterns 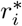 that are left invariant by a step of the RNN dynamics, in the absence of velocity inputs

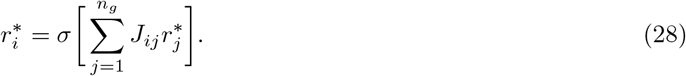

We identified slow points of the network dynamics by minimizing the scalar function

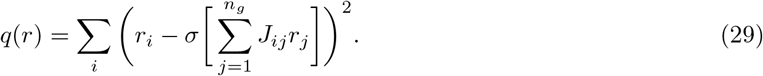

using the procedure outlined in [47]. We collected a set of 100^2^ fixed points by initializing the network on a grid of 100 × 100 spatial locations in the environment.

In order to identify the low-dimensional structure of population activity, we computed three spatial phases for each neuron’s rate map,

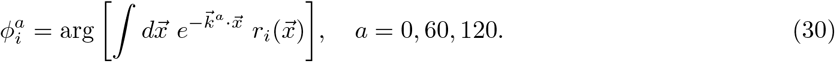

where 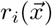 is the activity of neuron *i* at location 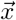, and 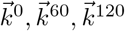 are the 0°, 60°, and 120° unit vectors. We then projected the population activity onto the following three pairs of axes,

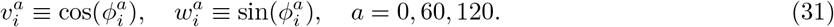

If each neuron in the network was a perfect hexagonal grid cell, then its firing rate could be written as,

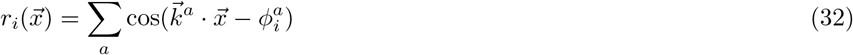

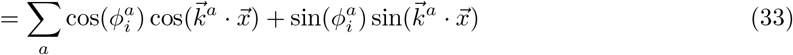

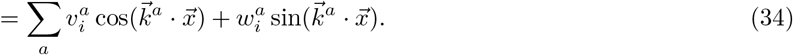

Hence projecting the population activity onto the three pairs of axes 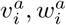 defined above would reveal a set of three perfect rings. In the case of the trained neural network, projecting the slow points onto the three pairs of axes 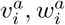 yields three pronounced rings 4D. The subspace spanned by the six vectors 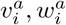 explains 52% of the total variance of the population activity.

As an additional measure to ensure that the twisted torus captures the dominant structure of the attractor manifold, we computed pairwise distances 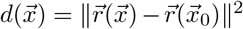 between the population activity 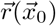 at a reference point in the environment, and the population activity 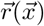 at all other points in the environment, and plotted 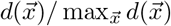 as a function of space 4C.

### 4.6 Idealized path integrator models

For all experiments probing trained RNN mechanisms, we implemented an idealized model for comparison (cf. sections 2.4–2.7 and Figures 5–8). Idealized models were designed to capture the two core mechanisms of previously published models: center-surround connectivity and offset recurrent weights. Update equations were

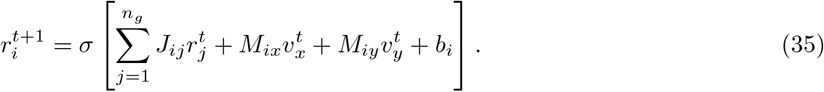

where *J* is the recurrent weight matrix, *M* is the velocity input weights, *v* is 2-dimensional velocity of the agent, *b* is a constant vector representing feedforward drive, and *σ* is the neuron nonlinearity. Neurons were placed uniformly over a grid of integer points on a 2-dimensional neural sheet with side length 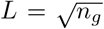. For two neurons with neural sheet positions 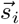 and 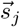, synapse weight *J_ij_* was set as

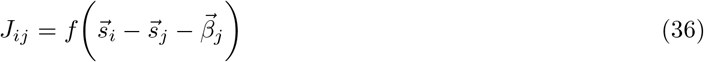

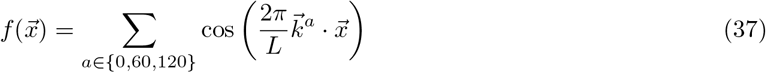

where 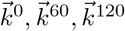 are the 0°, 60°, and 120° unit vectors as defined above and *β_j_* is the outgoing connectivity bias of neuron *j* For all models, we set *b_i_* = 1. Following [11], for a neuron with neural sheet position 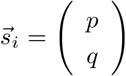 we set *M_ix_* = (*q* mod 2) (−1)^*p*^ and *M_iy_* = (*p* mod 2) (−1)^*q*^ (equivalent to dividing the neural sheet into 2×2 neuron zones each with a north, east, south and west-motion sensitive cell).

For Figure 7, 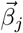 was set to 0 (ie. weights had no offset) to isolate the stable firing mechanism. For later figures probing updating mechanisms, 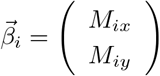 (intuitively, cells recieving north (south) velocity input have north (south) connectivity offsets; similarly for east/west). For Figure 8, the nonlinearity *σ* is the rectifying linear function.

The idealized 1-dimensional path integrator model referenced in Figures 5, 6 was implemented analogously. The update equation was the same as above, with 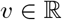 now representing the agent’s 1-dimensional velocity. Each neuron now has a position *s_i_* in a 1-dimensional neural ring. The weight between neurons at ring positions *s_i_* and *s_j_* was set analogously as

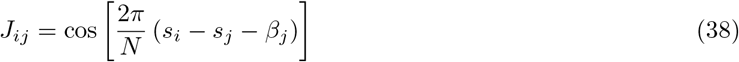

For all models, we set *M_i_* = (−1)^*i*^ and *b_i_* = 1. For Figure 5, *β_j_* was set to 0 (ie. weights had no offset) to isolate the stable firing mechanism. For later figures probing updating mechanisms, *β_i_* = (−1)^*i*^ *δ*. For Figure 6, the nonlinearity *σ* is the rectifying linear function.

### 4.7 Sorting RNN units onto a neural line and a neural sheet

#### 4.7.1 1-dimensional path integrator networks

Two methods were employed for sorting units in 1-dimensional path integrator networks onto a neural line, one based on neuron activity and one based on neuron connectivity. The first method consists of sorting neurons by preferred head direction (defined as the angle eliciting peak response for each neuron). This sort was used to produce the Fourier-transformed weight matrix in Figure 5F and for Figure 6.

To sort neurons by connectivity, each neuron *i* was assigned a random embedding coordinate *θ_i_*. The *θ*’s were then adjusted to maximize the energy function 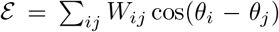 by gradient ascent in the direction: 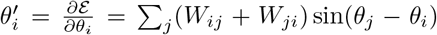. The dynamics were simulated until improvement in 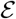 was small. The resulting *θ* coordinates represent a continuous approximation to the discrete neuron ordering that optimizes the similarity between the weights and the center-surround connectivity matrix 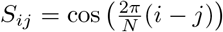. Neurons were then ordered by their *θ* coordinate, giving the final sort. This sort was used for Figure 5H.

#### 4.7.2 2-dimensional path integrator networks

The guiding intuition behind sorting was that network mechanisms would be easier to discern if neurons with similar spatial response properties were physically nearby on the neural sheet (Figures 7, 8). To do this, we obtain rate maps for all units in the network. For each neuron’s map, we compute three spatial phases

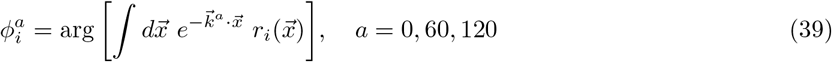

where 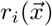 is the activity of neuron *i* when the model is at spatial location 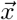, and 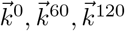 as above are the 0°, 60°, and 120° unit vectors. In the case of a perfect grid map, these phases uniquely determine the rate map of the neuron to be

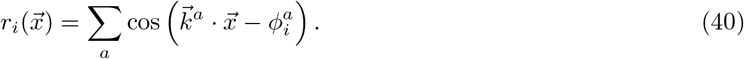

Under an ideal sort, moving from neuron to neuron along the neural sheet should continuously shift the rate map. We’ll posit that this shift of the rate map is proportional to the displacement on the neural sheet - that is a grid cell at neural sheet location 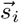 would have rate map

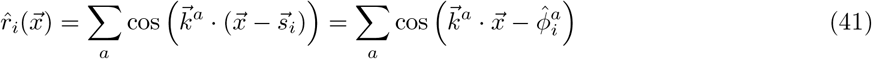

where 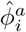 are the “ideal phases” associated with a neuron at sheet location 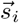 (and where we’ve slightly abused notation by treating 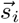, which is a neural sheet location, as a real-space location in the the formula above. In reality, there would be a proportionality constant relating neural sheet units to real-space units, which we take to be equal to 1 and ignore here).

To approximate this ideal sort, we optimize the position of neuron *i* on the neural sheet to match the ideal phases 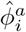 to the measured phases 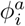:

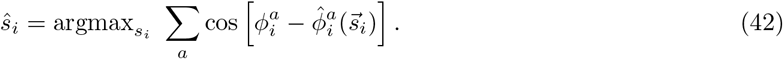

This gives a set of 2d coordinates for each neuron 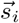. We bin the first coordinate into quantiles (*n* = 64) to obtain each neuron’s first sheet index, and then sort the neurons in each quantile by their second coordinate to obtain each neuron’s second sheet index (this procedure can be thought of as discretizing the first coordinate and then performing a lexographic sort on the second coordinates). The resulting indices place each of the 4096 neurons onto a single, unique node within a 64×64 grid. This 2D sorting procedure was used to extract the neural sheet from the activity maps in Figures 7 and 8.

### 4.8 Fourier analysis of recurrent weights

We used Fourier analysis on the connectomes of our trained networks to explain the stability of their spatial representations (sections 2.4, 2.6, Figures 5, 7). 1-dimensional network units were sorted by preferred head direction. The weight matrix was reordered to reflect this sort. The matrix was then transformed to the real Fourier basis using the usual change of basis formula, 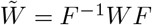, where *F* is the real Fourier matrix defined as

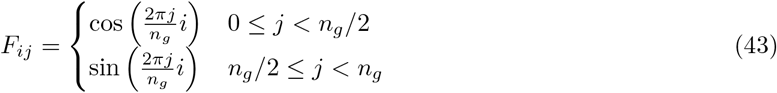

with appropriate normalization of each column. For the plots in Figure 5C,F, the Fourier transformed weight matrices were shifted so that low frequency pattern weights appear in the middle of the weight matrix (“fftshift”), and the matrix was cropped to a smaller window around the center so peaks were more easily visible.

For 2-dimensional networks (section 2.6), units and weight matrix were first sorted by map phases as described above. The weight matrix was then reshaped to have 4 indices - the 2-d neural sheet coordinates of the input, (*j_x_, j_y_*), and output (*i_x_, i_y_*) neuron. The matrix was then transformed to the 2d Fourier basis, now using the 2d Fourier matrix defined as

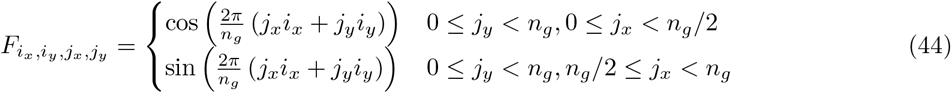

again with appropriate normalization of each 2-dimensional column 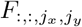. Change of basis was accomplished by the analog of the usual change of basis formula 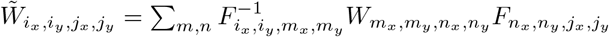. For plots in Figure 7C,G, we again shifted low frequencies to the center, extracted the diagonal 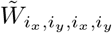 and cropped around the center.

#### Peak strength statistics

To quantify the significance of weight matrix “peaks” in Fourier space (cf. Figure 6) we defined the peak strength as the fraction of matrix power on the lowest frequency sine and cosine self-weight: 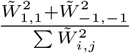. We trained 1000 1-dimensional path integrator networks using the protocol above, and obtained initial and final weight matrices under initial and final sorts (initial and final sorts defined below). These scores are histogrammed in 3.

Two null distributions are shown in Supplementary Figure 3. The first is the distribution of peak strengths of randomly generated matrices (entries drawn iid from a standard normal distribution) in their original sort (Supplementary Figure 3B, first column). The second is the distribution of peak strengths of optimally sorted random matrices (Supplementary Figure 3B, second column). Matrices were generated as before, but were then sorted by the connectivity-based sort designed to maximize peak strength described above (Sec. 4.7).

### 4.9 Linearized RNN dynamics

To gain a more quantitative understanding of how the 4 circuit components described above interact, we use the RNN’s update equations to track the effect of a small velocity input *dv* added to a stable bump pattern (see Figures 6, 8 and sections 2.5, 2.7).

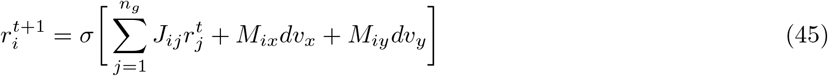

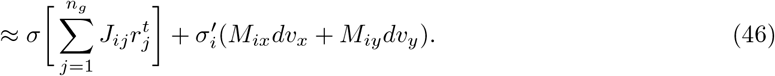

where *σ*′ is a diagonal matrix containing the derivatives of the neuron nonlinearities. Because the system is assumed to be at a fixed point, we can replace 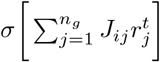 with 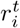, giving

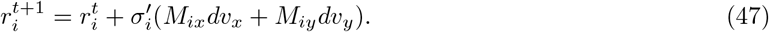

The term on the right captures the effect of the velocity cell activation *dv*, travelling through its input weights *M*, and subsequently through the neuron nonlinearity *σ*′. Note, however, that after one step, the velocity input has not yet passed through the offset recurrent weights *J*. This will only occur during the *next* step:

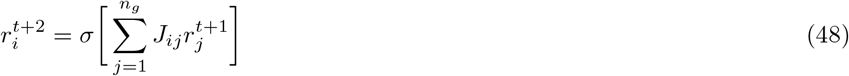

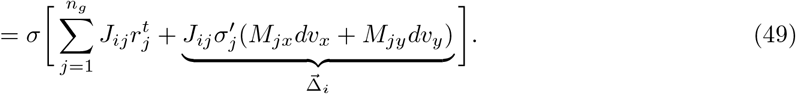

To compute 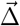, we note that because *σ* is the rectifying linear function, 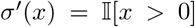, so that 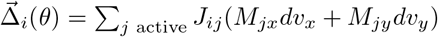.

For the 1-dimensional network (section 2.5 and Figure 6), the ground truth shifting term 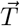 is defined as the tangent to the attractor manifold, 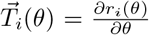, where *r_i_*(*θ*) is the activity of the *i^th^* neuron for head direction *θ*. This can be approximated as the difference between patterns at two nearby head directions: 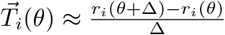.

For the 2-dimensional network (section 2.7 and Figure 8), the empirical and ground truth shifting terms depend on the 2-dimensional velocity 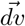. The same analysis as above gives 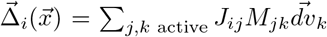 for the empirical shifting term. The ground truth shifting term was computed as 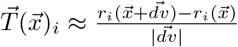, or the partial derivative of the attractor manifold in the direction 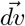.

### 4.10 Extracting Fourier modes from top connectivity eigenvectors

#### 2d RNN

Just as the population activity structure was not clear when projected onto its top PC eigenvectors, but became clear when we rotated to a more useful subspace, the top eigenvectors of the connectivity matrix *J* appear as linear combinations of low-frequency plane waves when viewed on the sorted neural sheet. However, with a simple orthogonal combination of the top eigenvectors, we can disentangle the consituent plane waves and construct 3 pairs of approximate pure modes, corresponding to the 3 pairs of plane waves used in traditional attractor model (section 2.6, Figure 7). The eigenvectors of the traditional attractor model can be written as

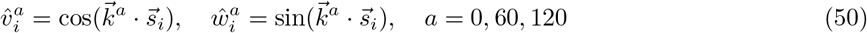

To see how closely the top eigenvectors 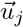, *j* = 1,…, 10, of the trained RNN connectivity matrix *J* approximate the perfect plane waves of the traditional model, we find the best orthogonal combination of the Uj. The connectivity matrix *J* of the trained RNN has ten large eigenvalues (Figure 7J), so we use only the top ten eigenvectors. Collecting the continuous attractor eigenvectors in *n_g_* × 6 matrix 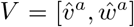, and the top ten trained network eigenvectors in a *n_g_* × 10 matrix 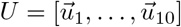, we identify the optimal linear transformation *O*, a 10 × 6 matrix, by minimizing the mean squared error,

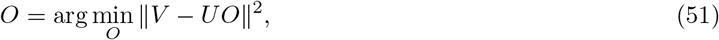

We then orthogonalize *O* using the Lowdin symmetric orthogonalization 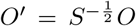 where *S* is the symmetric overlap matrix *S_ij_* = *O_i_ · O_j_*, and 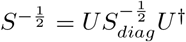, where *S_diag_* is obtained by diagonalizing *S, S_diag_* = *U*^†^*SU*. The transformed connectivity eigenvectors 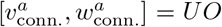 look like pure plane waves across the neural sheet, closely matching those in the traditional attractor model (Figure 7H).

### 4.11 Connectivity bias

In idealized path integrator networks, pattern updating is accomplished by discrete groups of cells with biased outgoing connectivity. To determine whether a similar pattern of biased outgoing connectivity exists in the trained networks, we first sorted the neurons onto a neural sheet (see Methods section 4.7 above). Then, for each neuron, we defined the connectivity bias as the displacement from the neuron’s own neural sheet position to the center of mass of its outgoing synaptic weights over the neural sheet:

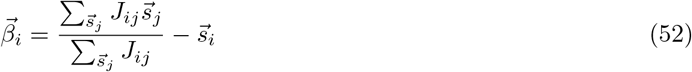

Defined this way, if a neuron projects isotropically around itself on the neural sheet, its connectivity bias is 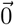.

The 2-dimensional displacements for the trained 2D integrator network are histogrammed in Figure 8E (right), along with those of the idealized model (left; showing one population of north, west, south, and east projecting cells).

## A Supplementary figures

**Supplementary Figure 1:**
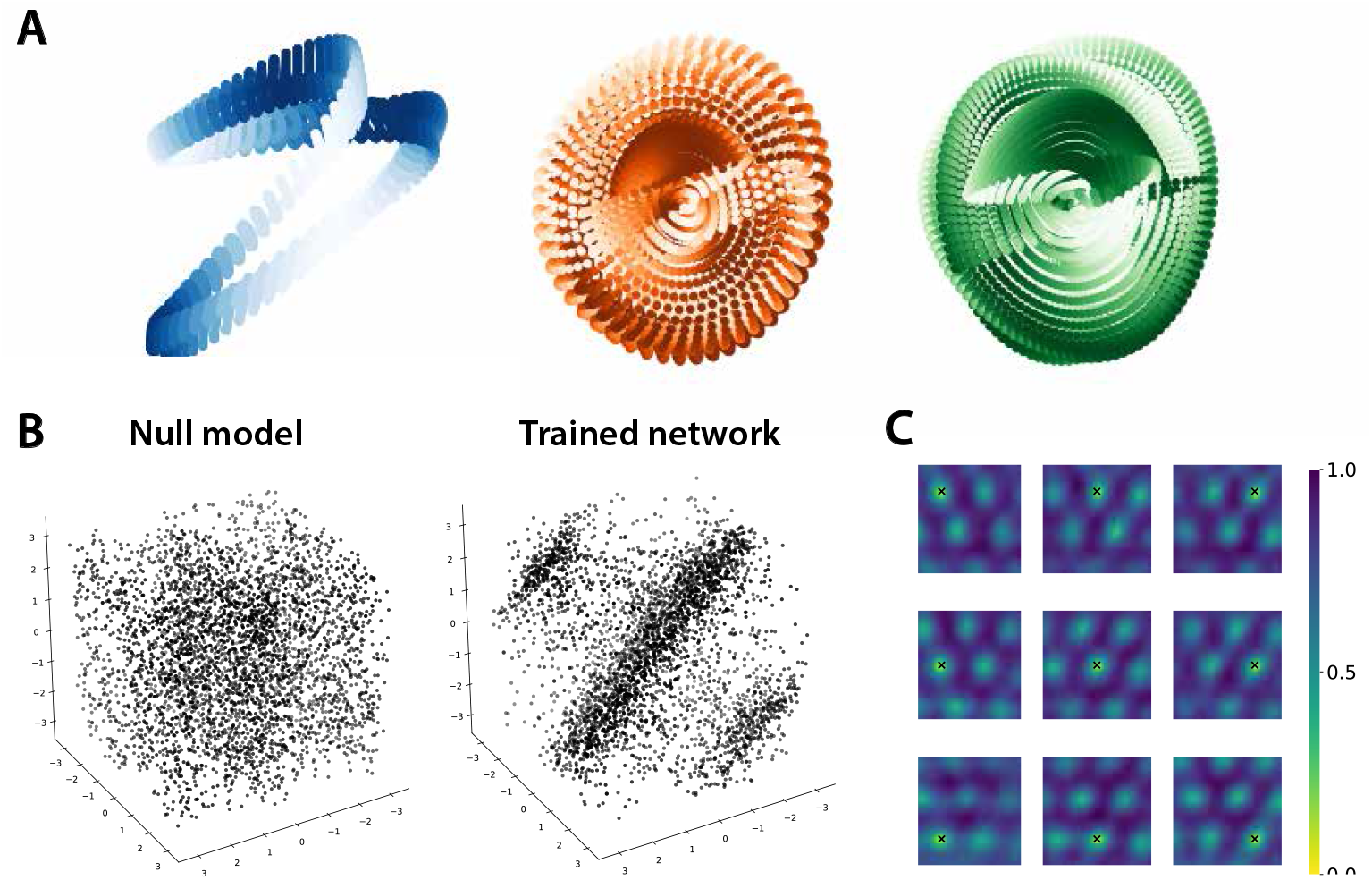
Additional attractor manifold analyses. Related to Figure 4. A) Ring-finding procedure applied to a null model of random low-passed noise patterns does not exhibit clear ring-like structure (see Methods 4.5 for details). B) left, This is due in part to the fact that the phases of the null model patterns do not obey a linear relationship. B) right, The phases of the grid patterns in the trained 2d integrator network, on the other hand, obey the relationship *ϕ*_1_ + *ϕ*_2_ + *ϕ*_3_ = 0. C) Distance between the activity pattern at a reference point in the 2d environment (marked with a black x) and the activity pattern at all other points in the 2d environment (activity pattern of top 25% grid-scored cells only; see Methods 4.3), plotted for several different reference points. When only the top 25% of grid cells are included, the hexagonal periodicity is much clearer than when all cells are included (Figure 4C), indicating that the remaining cells introduce substantial heterogeneity.

**Supplementary Figure 2:**
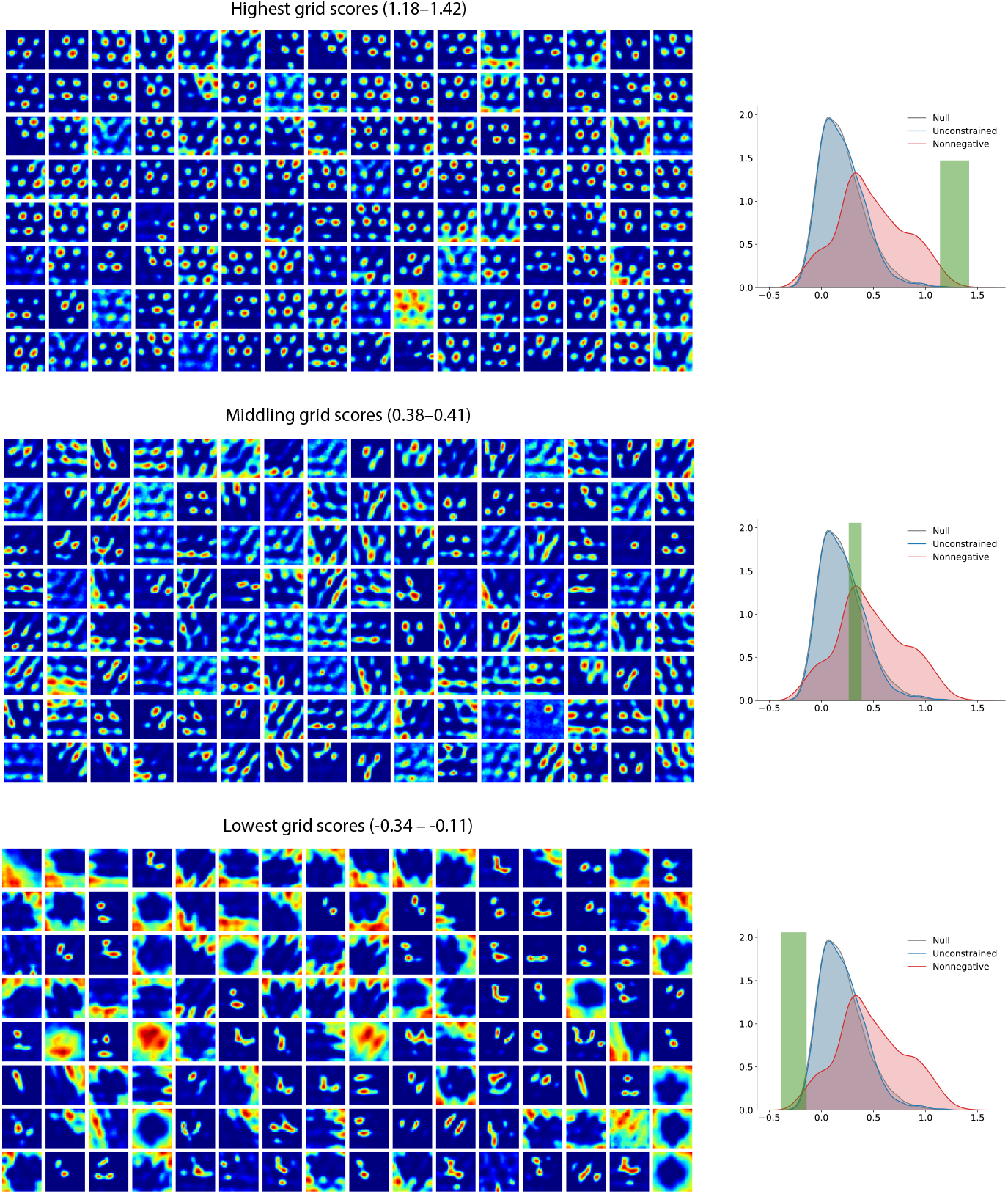
A selection of grid cells from the trained network, sorted by grid score. Related to Figure 4. Rate maps taken from the trained 2d integrator network shown in Figure 4. Top panel, 128 neurons with highest grid score (left), with grid scores in the interval shown in green (right). Middle, 128 neurons (left) taken from the middle of the grid score distribution (green window, right). All 128 are above the grid score threshold, 0.37, used to define a grid cell in [31]. Bottom, 128 neurons with lowest grid score (left), with grid scores in the interval shown in green (Right). These cells tend to be quite varied, and many resemble border cells, spatial non-grid cells, and other heterogeneous types described in medial entorhinal cortex. See Methods 4.3 for details on how grid score was computed.

**Supplementary Figure 3:**
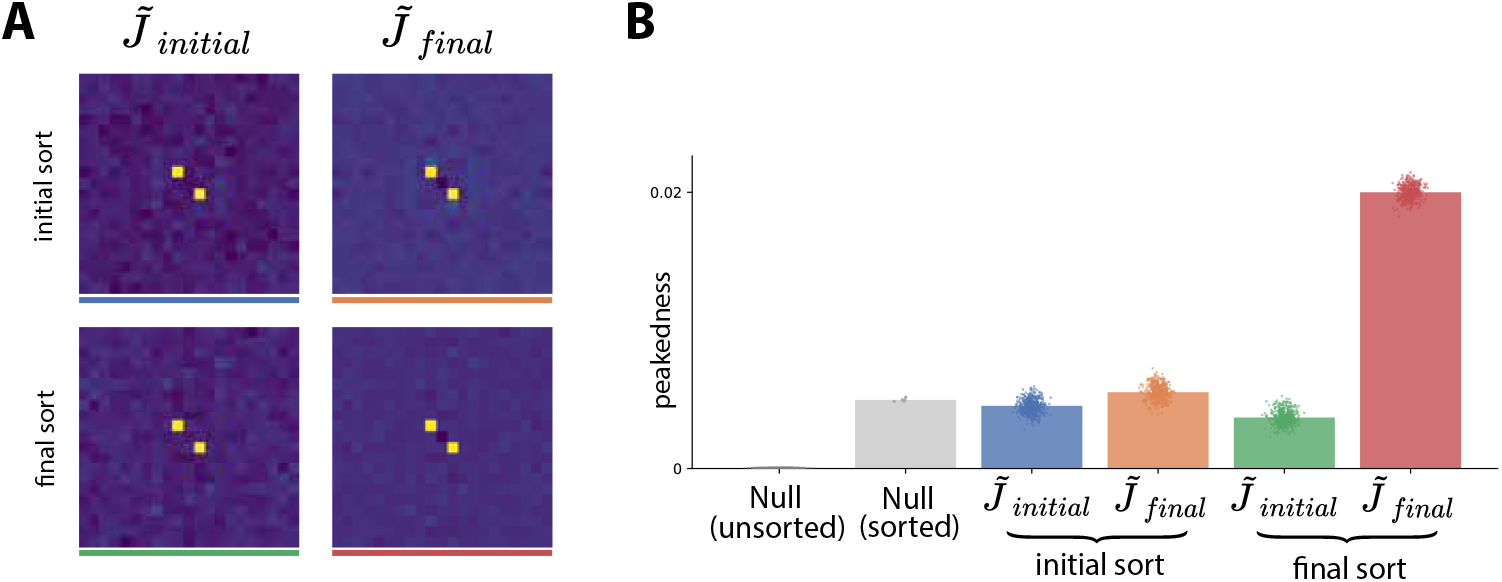
Training discovers and amplifies latent attractor dynamics in random matrices. Related to Figure 6. A) The 1D path integrator network’s weight matrix in Fourier space before and after training, under two different sorts. Left column, the weight matrix at random initialization prior to any training. Right column, the weight matrix after full training. Top row, optimal sort for initial pretraining matrix. Bottom row, optimal sort for final post-training matrix (see Methods 4.7 for details on how sort was obtained). Under the final sort (bottom row), as we show in Figure 5, the trained weights are highly peaked in Fourier space (bottom right). However, when the neurons are arranged this way, the initial weights appear similarly peaked (bottom left), which is surprising given that it is a randomly initialized matrix with no training. B) Plotting the Fourier peakedness (see Methods 4.8 for details) for the matrices clarifies these results. Under the final sort, the initial weight matrix shows peakedness similar to a random matrix under its best possible sort (compare green and grey bars), and much higher than an unsorted random matrix (first bar, “Null (unsorted)”). This suggests that, given a random initial weight matrix, training discovers a near-optimal sort corresponding to a rough attractor network hidden in the random weights (the final sort, green bar), and over time accentuates this structure by increasing Fourier peakedness (red bar).

**Supplementary Figure/Video 4:**
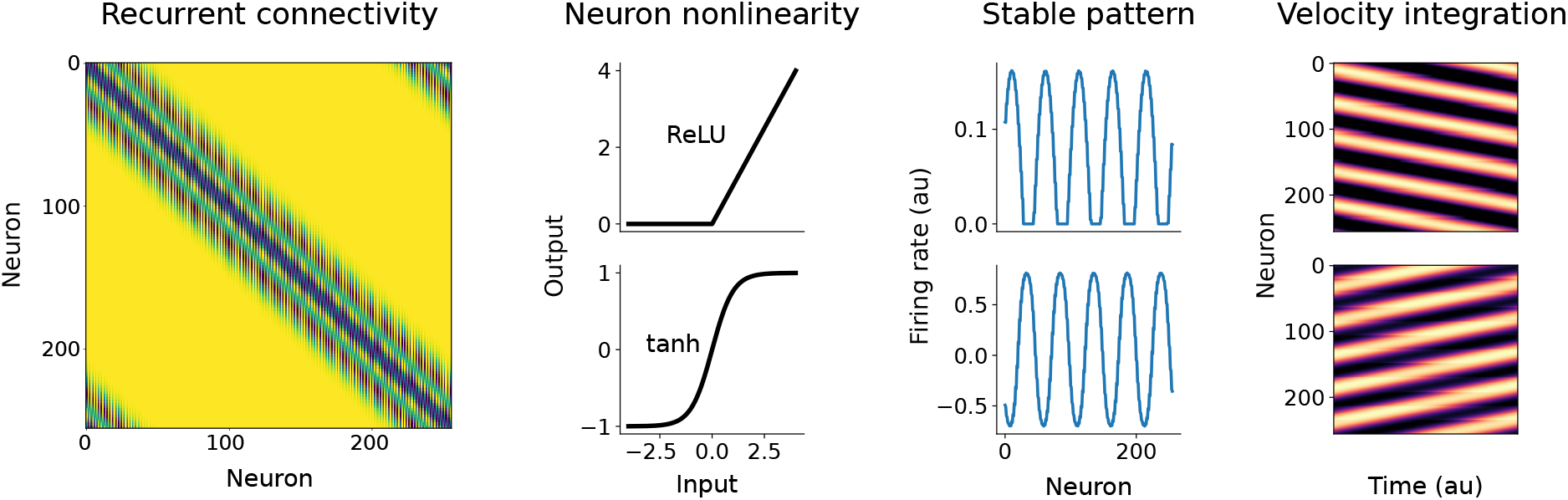
Connectivity alone is not enough to determine the behavior of a path integrator circuit. Related to Figure 6. Frame taken from Supplementary Video. We created two parallel 1d path integrator models (see Methods 4.6 for details). Left, the recurrent connectivity and velocity input weights were identical for the two models. Cells connect through short range excitation and long-range inhibition, and each cell has a a specific systematic offset in its outgoing connectivity, either left or right (see Methods 4.6). Center left, in one model, we chose a rectifying neuron nonlinearity (top), while in the other, we chose tanh (bottom). Center right, the resulting stable activity patterns are qualitatively very similar, and consist of localized clusters of active neurons interleaved with clusters of inactive neurons. Right, we provided additive input to cells with rightward-shifted outgoing connectivity and examined the resulting recurrent dynamics over time. Video shows the resulting population activity as a time-varying curve. Here, we show neuron activities over the whole population, over the whole stimulation period, as a heatmap. Vertical slices show the full population activity at an instant in time, and the horizontal axis shows the change in population activity pattern over time. In the model with a rectifying nonlinearity, periodic patterns continuously shift rightward, as expected when stimulating right-shifting cells. On the other hand, the model with the tanh nonlinearity integrates in the opposite direction. This highlights the important of the term corresponding to the nonlinearity in our change term analysis (see section 2.5 for overview and Methods 4.9 for mathematical details).

## Footnotes

1 Note that cc. is shorthand for complex conjugate. For any real solution *g*(*x*) to Eq. 21, 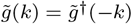. Therefore, for each wavevector *k* we must also include its negative, −*k*.

